# Systematic benchmarking of basecalling models for RNA modification detection with highly-multiplexed nanopore sequencing

**DOI:** 10.1101/2025.07.11.663424

**Authors:** Gregor Diensthuber, Ivan Milenkovic, Laia Llovera, Ana Milovanovic, Francesco Pelizzari, Eva Maria Novoa

## Abstract

Nanopore direct RNA sequencing (DRS) holds promise for advancing our understanding of the epitranscriptome by detecting RNA modifications in native RNA molecules. Recently, Oxford Nanopore Technologies (ONT) has released basecalling models capable of detecting several RNA modifications. However, their accuracy, sensitivity and specificity, as well as cross-reactivity against other modification types, remains largely unexplored. Here, we systematically benchmark modification-aware models by evaluating their performance on a highly-multiplexed panel of synthetic molecules covering all possible sequence contexts, as well as on biological samples from a diverse set of species. We find that modification-aware models reliably detect diverse RNA modification types across a broad range of sequence contexts. However, they are prone to elevated false positive rates and exhibit notable cross-reactivity with other RNA modification types. We show that the use of modification-free controls allows significant, yet incomplete removal of false positives, thus constituting an essential control. Finally, we demonstrate that basecalling error patterns and alterations in current features can identify differentially modified sites, for modifications for which modification-aware models are absent. Overall, our results underscore the utility and accuracy of modification-aware basecalling models for RNA modification detection, while highlighting the importance of including diverse control samples to mitigate false positive rates.

## INTRODUCTION

RNA modifications, collectively referred to as ‘the epitranscriptome’, are chemical moieties present in RNA molecules, fine-tuning their structure, function and molecular fate (1, 2). In recent years, the possibility to sequence native RNA molecules has emerged, enabling the direct capture of RNA modifications without the need for reverse transcription or PCR amplification, bypassing the need for selective antibodies or chemical probing reagents (3–5). This has been enabled by the nanopore direct RNA sequencing (DRS) platform, currently commercialised by Oxford Nanopore Technologies (ONT) (6).

To detect RNA modifications, approaches developed so far can be binned into three main categories: methods that rely on systematic basecalling errors, which exploit the fact that RNA modifications cannot be base-called by models that were only trained to detect canonical bases (A, C, G, U) (7–12); i) methods that rely on alterations in raw current features, such as signal intensity, trace or dwell time (13–20); and iii) modification-aware basecalling models, which have been trained to detect a modification of interest, and thus can directly predict RNA modifications in individual reads, in addition to the four canonical bases (21). Recently, ONT has released several modification-aware basecalling models, namely for *N6*-methyladenosine (m^6^A), inosine (I), 5-methylcytosine (m^5^C) and pseudouridine (Ψ). These models offer the possibility to predict RNA modifications *de novo,* with single nucleotide and single molecule resolution. However, their accuracy, false positive rates and cross-reactivity with other RNA modifications remain unclear.

Here we assess the performance of modification-aware basecalling models released (September 2024) by ONT (see **Table S1**) on synthetic modified and unmodified RNA molecules, as well as on rRNAs from diverse species (**Table S2**). Globally, our results show that modification-aware basecalling models capture the most modified sites at high stoichiometries, but can mispredict a significant number of sites. Moreover, some models show strong cross-reactivity with other RNA modifications found on the same nucleobase, which could lead to inaccurate conclusions if the models are used for *de novo* prediction of RNA modifications. We find that the use of modification-free controls that cover equivalent sequence contexts to the samples of interest (12, 22) are necessary to remove false positives from nanopore basecalling models. However, we also show that these controls are insufficient to remove all false positives, warranting future improvements to the basecalling models or correction methods. Finally, for those RNA modifications for which modification-aware basecalling models are not available, we demonstrate that diverse RNA modification types can be identified in the form of differential base-calling errors (9, 10, 23) and altered current features, demonstrating that previously developed approaches for RNA002 chemistry can still be used in the context of the new RNA004 chemistry.

## RESULTS

### Expanding direct RNA-sequencing demultiplexing to 96 barcodes for RNA004 chemistry

To ease the systematic benchmarking of diverse synthetic and *in vivo* RNA molecules with known modifications, we reasoned that a multiplexing model to barcode all samples in a single flowcell would greatly simplify the experimental setup and downstream bioinformatic analysis, as well as significantly minimize the batch effects observed across samples that are sequenced in different flowcells. However, no commercial barcoding kit exists for the latest direct RNA sequencing chemistry (SQK-RNA004), limiting the technology’s use and cost-effectiveness. To this end, community-driven efforts have recently been released for multiplexing DRS runs (24–26), some of which are compatible with the newest chemistry, showing high precision and recall (25, 26). However, these efforts are so far limited to multiplexing a few barcodes in the case of RNA004 chemistry.

Here, we extended previous efforts by training a model capable of demultiplexing up to 96 samples with the latest SQK-RNA004 kits (see **Table S3** and *Methods*). We found that the newly trained 96-barcoding model (b96_RNA004) reached a precision of 99.3% with 100% recall on the validation dataset (**Fig. S1A**). Furthermore, the processing time was not significantly affected compared to our four-barcode model (b04_RNA004), resulting in a mean real-time of 1.3 min per 100,000 reads processed (**Fig. S1B,** see also **Table S4**). Finally, we tested the performance of the b96_RNA004 demultiplexing model on an additional independent test dataset in which barcodes 21, 59, and 88 were absent from the run (*see Methods*). We found that 0.00009% of the total reads were misassigned to either of these barcodes after the default cutoff of baseQ > 50 was applied (**Fig. S1C**), suggesting that this model provides a highly accurate and rapid way to sequence up to 96 samples in a single flowcell.

### Systematic benchmarking of modification-aware RNA basecalling models with synthetic sequences

In September 2024, Oxford Nanopore Technologies (ONT) released a set of modification-aware basecalling models (m6A_DRACH, inosine_m6A, pseU, and m5C) that can detect four different RNA modification types (m^6^A, I, Ψ and m^5^C) using their proprietary basecalling software *dorado* (https://github.com/nanoporetech/dorado). To systematically benchmark these models on datasets with known ground-truths in which the modification stoichiometry at each position is known, we employed synthetic RNA constructs covering all possible 5-mer sequences (n = 1024), referred to as ‘curlcakes’ (7), which have been previously used to assess the detection of diverse RNA modification types using the DRS platform (16, 27), as well as to train diverse algorithms to detect RNA modifications (16, 19, 28, 29). Specifically, here we generated ‘curlcake’ constructs with either canonical nucleobases (A, C, G or U) –the unmodified control–, or with one different modified nucleobase at a time, including those for which modification-aware basecalling models have been trained (m^6^A, m^5^C and Ψ), and for other RNA modification types for which there are currently no modification-aware basecalling model available (hm^5^C, ac^4^C, m^1^Ψ and m^5^U). Next, highly-multiplexed DRS libraries of synthetic modified and unmodified controls were generated in triplicates and sequenced on a PromethION flowcell (FLO-PRO004RA) (**Fig. 1A**). Following sequencing, the raw sequencing data was basecalled with either of the four modification-aware basecalling models using the MasterOfPores nextflow workflow (**Fig. S2A-B**) (30). Finally, per-read modification probabilities were extracted using the ONT *modkit* software (https://github.com/nanoporetech/modkit), and per-site modification stoichiometries were calculated using default modification probability thresholds (**Fig. 1B**) (see also *Methods*).

**Figure 1.**
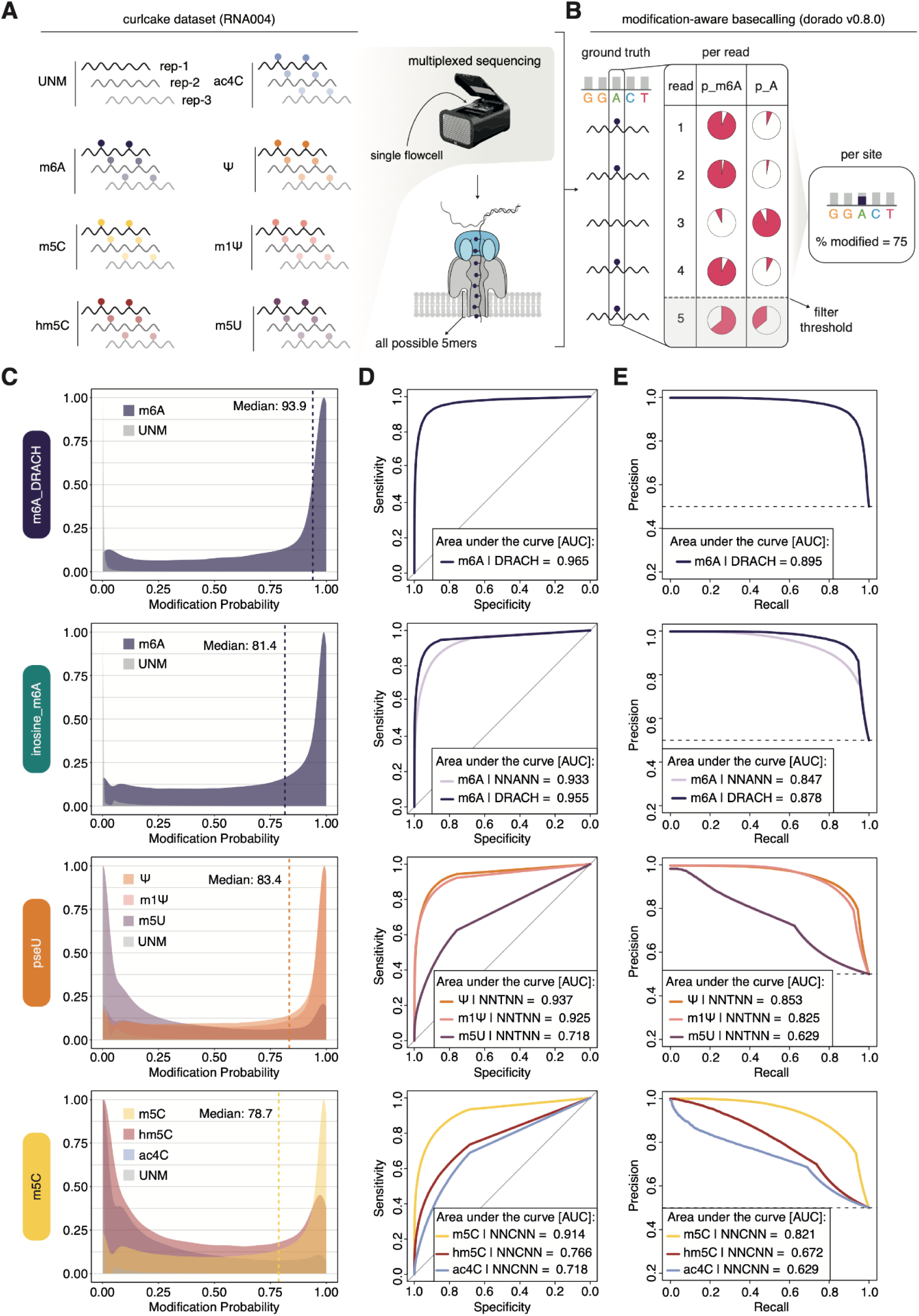
Systematic assessment of the accuracy and cross-reactivity of pre-trained modification-aware basecalling models across a panel of synthetic sequences with diverse RNA modifications. **(A)** Schematic representation of the experimental setup used to systematically benchmark modification-aware basecalling models. Synthetic ‘curlcake’ constructs, which contain all possible 5-mer combinations (n = 1024), were generated for a subset of RNA modifications (m^6^A, m^5^C, hm^5^C, ac^4^C, Ψ, m^1^Ψ, and m^5^U), including their unmodified counterpart (UNM) and sequenced with the SQK-RNA004 kit in a highly-multiplexed fashion using a single flowcell. All constructs were basecalled using *dorado* modification-aware basecalling models (m6A_DRACH, inosine_m6A, pseU and m5C). **(B)** Schematic representation of the information provided by modification-aware basecalling models (e.g. single modification m6A-aware model). First, the model assigns a probability for each read and position of either being modified (e.g. p_m6A) or unmodified (e.g. p_A). Subsequently this information can be collapsed into a per-site prediction using *modkit*, considering a threshold that is calculated per sample (default = 10^th^ percentile). Positions that are below the threshold are excluded from per site analysis (e.g. p_m6A and p_A of read 5 are below the threshold and therefore excluded). **(C)** Per-read modification probabilities reported for each model on fully modified and unmodified sequences. The vertical dashed line represents the median predicted modified frequency (expected = 100%). **(D)** ROC curves based on modification probabilities per read and site for each model. **(E)** Precision-Recall (PR) curve based on modification probabilities per read and site, for each basecalling model.

We first evaluated the performance of each model in identifying their designated RNA modification (e.g., m^6^A), compared to their unmodified counterpart (e.g., A). In the case of m^6^A detection, both the ‘all-context’ model (inosine_m6A) and the ‘context-aware’ model (m6A_DRACH) were examined. We should note that the latter restricts its predictions to the mammalian m^6^A modification consensus motif DRACH. We then extracted per-read modification probabilities across all possible 5-mers, finding that the models globally yielded median modification probabilities ranging from 0.79 (m5C model) to 0.94 (m6A_DRACH model). Thus, performance varied strongly depending on the model, with the m5C model being the one reporting the overall lowest modification probabilities at modified sites, as well as showing a higher overlap between the distribution of modification probabilities of modified and unmodified sites (**Fig. 1C**).

To quantitatively assess these differences, we then calculated the performance of each model at per-read level, and built Receiver Operating Characteristic (ROC) curves for each model (**Fig. 1D**). All models showed a good performance globally (ROC-AUC = 0.914-0.965), with the m5C model being the one showing lowest Area Under the Curve (AUC) values. Considering that m^6^A mainly occurs in DRACH sites, we then compared the performance of m6A_DRACH and inosine_m6A models to detect m^6^A modifications in DRACH sequence contexts, finding that the m6A_DRACH model showed improved performance (ROC-AUC=0.965) compared to the ‘all-context’ inosine_m6A) counterpart (ROC-AUC=0.955) (**Fig. 1D**). Similarly, Precision-Recall (PR) curves were built for each model to examine how the models would identify positive instances (true positives) while minimizing the false positives (**Fig. 1E**). Our analyses revealed that the AUC values ranged from 0.821 (m5C model) to 0.895 (m6A_DRACH model), suggesting that the former has a higher amount of false positives.

### Modification-aware basecalling models show cross-reactivity against other RNA modification types

Current ONT modification-aware basecalling models are trained to specifically identify a given modified base against its canonical counterpart (e.g., m^6^A versus A). However, whether these models cross-react with other similar chemical marks that also exist on the same nucleobase remains largely unexplored. To investigate the potential cross-reactivity of these models, we examined their performance on m^1^Ψ- and m^5^U-modified sequences (for the pseU model), and on hm^5^C- and ac^4^C-modified sequences (for the m5C model). This analysis revealed substantial cross-reactivity of both models against other modified bases (**Fig. 1C-E**). Indeed, the pseU model strongly cross-reacted with m^1^Ψ modifications (AUC = 0.925, PR-AUC = 0.853), with near-identical AUC values to those obtained against the primary target, pseudouridine (AUC = 0.937, PR-AUC = 0.825), suggesting that this model cannot distinguish the two modification types. Of note, for most *in vivo* analyses, this should still be tolerable, as *N^1^*-methylpseudouridine is a lowly abundant RNA modification occurring on few RNA biotypes, such as archaeal transfer RNAs (31). Additionally, this cross-reactivity could be exploited for examining the quality of mRNA vaccine candidates, which are largely generated using m^1^Ψ modifications (32), without the need of m^1^Ψ-specific models. By contrast, m^5^U cross-reacted to a lesser extent, yet still significantly better than random (AUC = 0.718, PR-AUC = 0.629). Similar results were observed when examining the cross-reactivity of the m5C model towards its hydroxylated version hm^5^C (AUC = 0.766, PR-AUC = 0.672) and ac^4^C (AUC = 0.718, PR-AUC = 0.629) (**Fig. 1D,E**, see also **Fig. S3**). We should note that in the case of m^5^C and hm^5^C, this cross-reactivity could potentially lead to misinterpretations on *in vivo* data, where the two modifications are known to carry out different functions (33). Notably, these results also suggest that the cross-reactivity of the base-calling models also significantly varied depending on the k-mer (**Fig. 1C**).

### K-mer specific biases of modification-aware basecalling models

Current ONT basecalling models have been trained with synthetic oligonucleotides in which the modified base is surrounded by a degenerate sequence, with the exception of the m6A_DRACH model. For this reason, they should in principle detect RNA modifications in all sequence contexts. However, previous works have shown that the alteration of the current signal relative to the unmodified counterpart varies depending on the sequence context (16, 34), suggesting that modifications embedded in certain k-mer contexts might be better predicted than others. To systematically examine the potential k-mer preference for detecting RNA modifications, we calculated the median modification probability for each 5-mer containing a single central modified base (n = 81), finding that certain kmers were likely underrepresented, reflected by lower median modification probabilities (**Fig. 2A**). Next, we performed motif enrichment analysis on those 5-mers whose modification probabilities were lower than 0.75 (*see Methods*), revealing that certain sequence contexts are indeed especially problematic for the detection of modifications. Notably, these contexts varied depending on the model; for example, A/U-rich contexts were troublesome for the m5C model, whereas GC-rich contexts showed decreased modification probabilities for detecting Ψ modifications (**Fig. 2B**).

**Figure 2.**
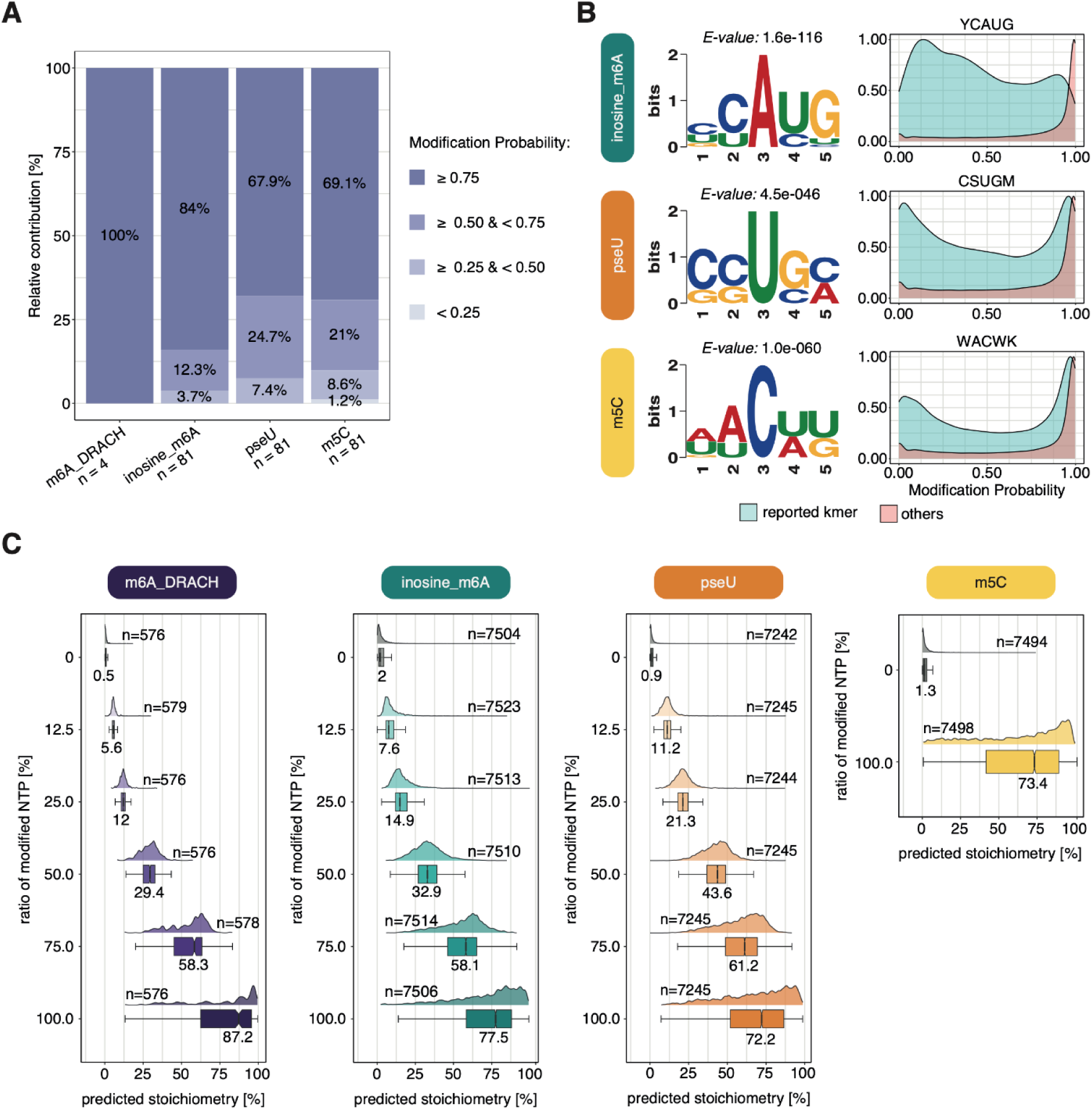
Kmer-specific biases of modification-aware basecalling models and stoichiometry estimates. **(A)** Barplot representing the median modification probability of single centrally-modified 5mers for the m6A_DRACH (KGACY, where K = G/T and Y = C/T, n = 4), inosine_m6A (BBABB, where B = T/C/G, n = 81), pseU (VVTVV, where V = A/C/G, n = 81) and m5C models (DDCDD, where D = A/G/T, n = 81). **(B)** Enriched motifs relative to the background distribution of kmers (*see Methods*) and corresponding density plots of modification probabilities for kmers with a median modification probability < 0.75. **(C)** Boxplots representing the predicted stoichiometry levels per position across a range of six modification levels (0%, 12.5%, 25.0%, 50.0%, 75.0% and 100.0% of modified NTP used in each *in vitro* transcription). Median values are indicated below each boxplot with *n* representing the sample size.

### Modification stoichiometry is underestimated by modification-aware basecalling models

Next, we examined the ability of modification-aware basecalling models to make correct predictions in terms of the relative amount of molecules that are modified at a given position, also referred to as percent modified or modification stoichiometry. To examine the accuracy of the models at predicting modification stoichiometry, we built ‘curlcake’ constructs that contained different proportions of modified NTPs (0, 12.5, 25, 50, 75 and 100%), and used the ONT *modkit* tool with default parameters to extract per-site modification levels (see *Methods*). The results revealed that all models systematically underestimated the stoichiometries present in the sample, with models predicting a median modification stoichiometry of 72.2-87.2% for samples that were 100% modified, and with high variance in modification stoichiometry across k-mers (**Fig. 2C**). We should note that the observed stoichiometry values will vary depending on whether the future user is only interested in a subset of k-mer sequence contexts.

### Benchmarking of modification-aware basecalling models on *in vivo* total RNA samples

We then evaluated the performance of the models on rRNA sequences from *E. coli* and *S.cerevisiae* for which accurate maps of rRNA modifications exist (35–37). Importantly, they contain a large number of Ψ modifications as well as several m^5^C modifications. For each species, total RNA was extracted and *in vitro* polyadenylated, prepared with standard DRS library preparation, and basecalled using either the pseU or m5C model (**Fig. 3A**).

**Figure 3.**
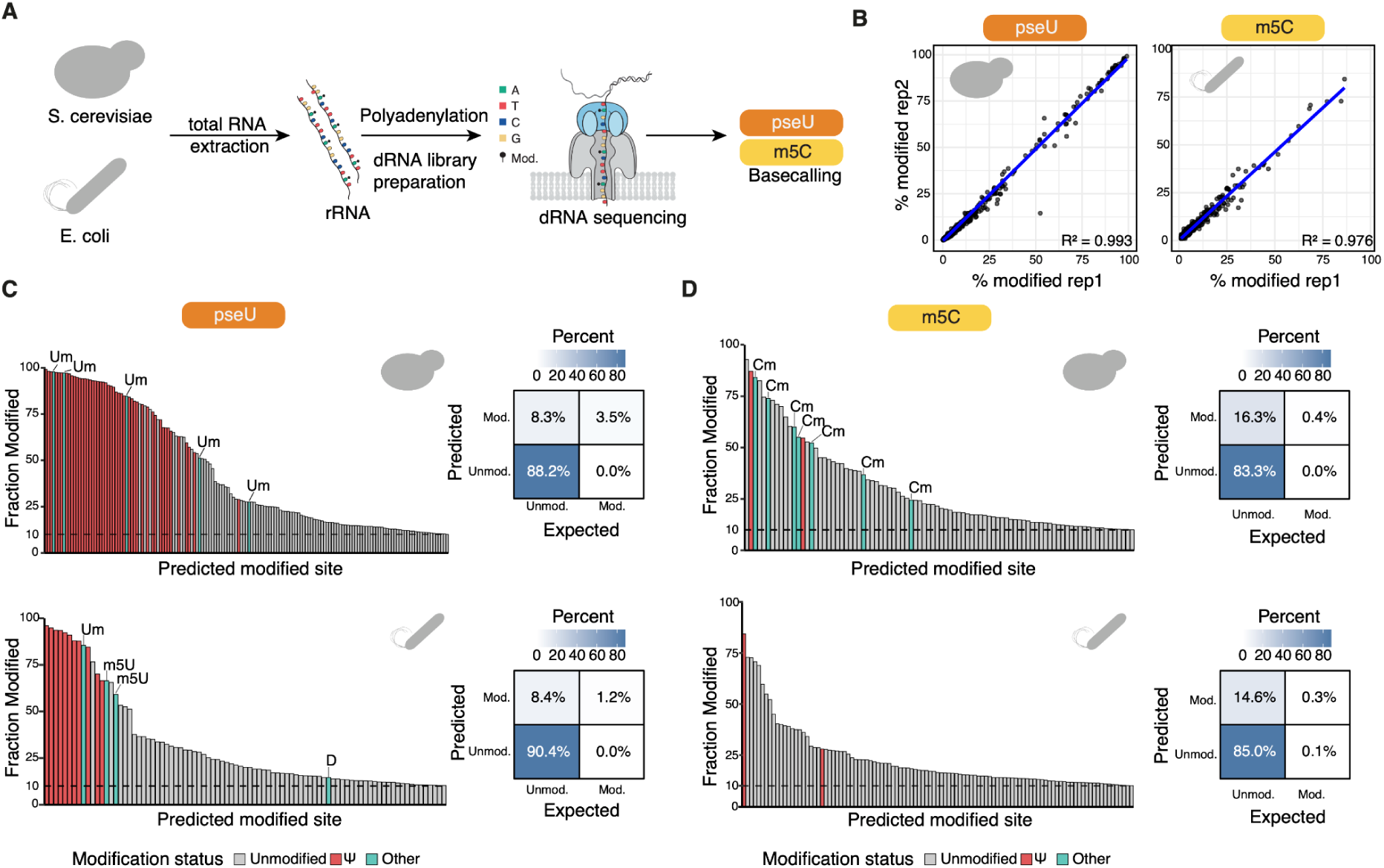
Performance of pseU and m^5^C modification-aware basecalling models on *S. cerevisiae* and *E. coli* rRNAs. **(A)** Schematic representation of the experimental setup to benchmark modification-aware models in in vivo samples. Total RNA from *S. cerevisiae* and *E. coli* was extracted and cleaned, in vitro polyadenylated, prepared for direct RNA sequencing using SQK-RNA004 kits. The samples were then basecalled using either N5-methylcytosine (m^5^C) or pseudouridine (pseU) modification-aware models (see *Methods*) in dorado. Modification information was extracted using the *modkit* software, developed by ONT. **(B)** Scatterplot depicting the predicted modification stoichiometry (% modified) using either the pseU or m5C models, in either *S. cerevisiae* or *E. coli* rRNA, respectively. **(C,D)** Histogram depicting the modification stoichiometry for each of the predicted modified sites in rRNAs from *S. cerevisiae* (upper panels) and E. coli (lower panels), basecalled with either pseU **(C)** or m5C models **(D)**, respectively. Predicted modified sites (by dorado+modkit) have been coloured depending on whether they are: i) true positive sites (red), ii) non-cognate modified sites (false positives) (blue), and iii) unmodified sites predicted as ‘modified’ (gray). A contingency table matching each histogram, depicting the relative proportion of false positives (mod-unmod), true positives (mod-mod), false negatives (unmod-mod) and true negatives (unmod-unmod), is also shown.

We first examined the replicability of the modification stoichiometry of predicted modified sites, finding that the estimates of modification stoichiometry were highly replicable across biological replicates in both species (R^2^=0.976-0.993) (**Fig. 3B**). However, we should note that a significant proportion of modified sites identified were lowly modified (<25%) albeit most sites are expected to be fully modified, based on previously published LC-MS/MS experiments (37–39).

We then examined whether the sites predicted by the pseU and m5C models overlapped with annotated Ψ and m^5^C sites, respectively (**Fig. 3C,D**). Indeed, we observed that a significant proportion of predicted Ψ sites (31% in yeast and 21% in *E. coli*) overlapped with annotated Ψ sites; however, several sites containing other modification types (e.g. Um and m^5^U) were also captured by the pseU model and thus were mispredicted as Ψ-modified sites (**Fig. 3C**) in addition to a significant proportion of unmodified sites (66% in yeast and 74% in *E. coli*). Similarly, the m5C model captured the two annotated m^5^C modified sites in yeast, and two out of three in *E. coli*, but also a vast proportion of false positives, which included both Cm-modified and unmodified sites (**Fig. 3D**). We should note, however, that most of the high-modification stoichiometry sites corresponded to true positives, and these were top-ranked in terms of modification stoichiometry, leading to very high AUC-ROC values (AUC-ROC:0.920-0.999), but more modest AUC-PR values (AUC-PR:0.21-0.90) (**Fig. S4**). Altogether, our results show that *in vivo*, modification-aware basecalling models are able to capture highly modified Ψ and m5C-modified sites, but at the expense of a substantial amount of false positives, including other modification types for which the models cross-react with (e.g., Um, m^5^U, Cm) as well as unmodified sites that are mispredicted as modified, suggesting that there is a strong need for external fully-unmodified controls to remove false positive predictions when using modification-aware base-calling models.

### Removal of false positives using modification-free synthetic matched controls

Previous works have proposed the use of modification-free control datasets to remove false positive predictions in epitranscriptomics (40–42). We therefore imagined that using such constructs would remove mispredicted sites reported by modification-aware basecalling models. To this end, we generated modification-free rRNA molecules from *E. coli, M. musculus,* and *H. sapiens* using *in vitro* transcription of cDNA products (IVT), previously amplified to incorporate a T7 promoter. Samples were then basecalled using either the pseU or m5C model to perform pairwise comparisons of the predicted per-site modification frequencies of the unmodified control samples (**Fig. 4A**, see also *Methods*). Our results revealed that a significant proportion of RNA modifications predicted in WT were also predicted in the IVT sequences, suggesting that IVT controls can be useful for the removal of false positive predictions (**Tables S5 and S6**). Notably, replicability of the predictions in the IVT sequences was very high (Pearson’s *r* = 0.88-0.99), suggesting that these false predictions are not random but rather systematic, and possibly related to the sequence context itself (**Fig. S5A-B**).

**Figure 4.**
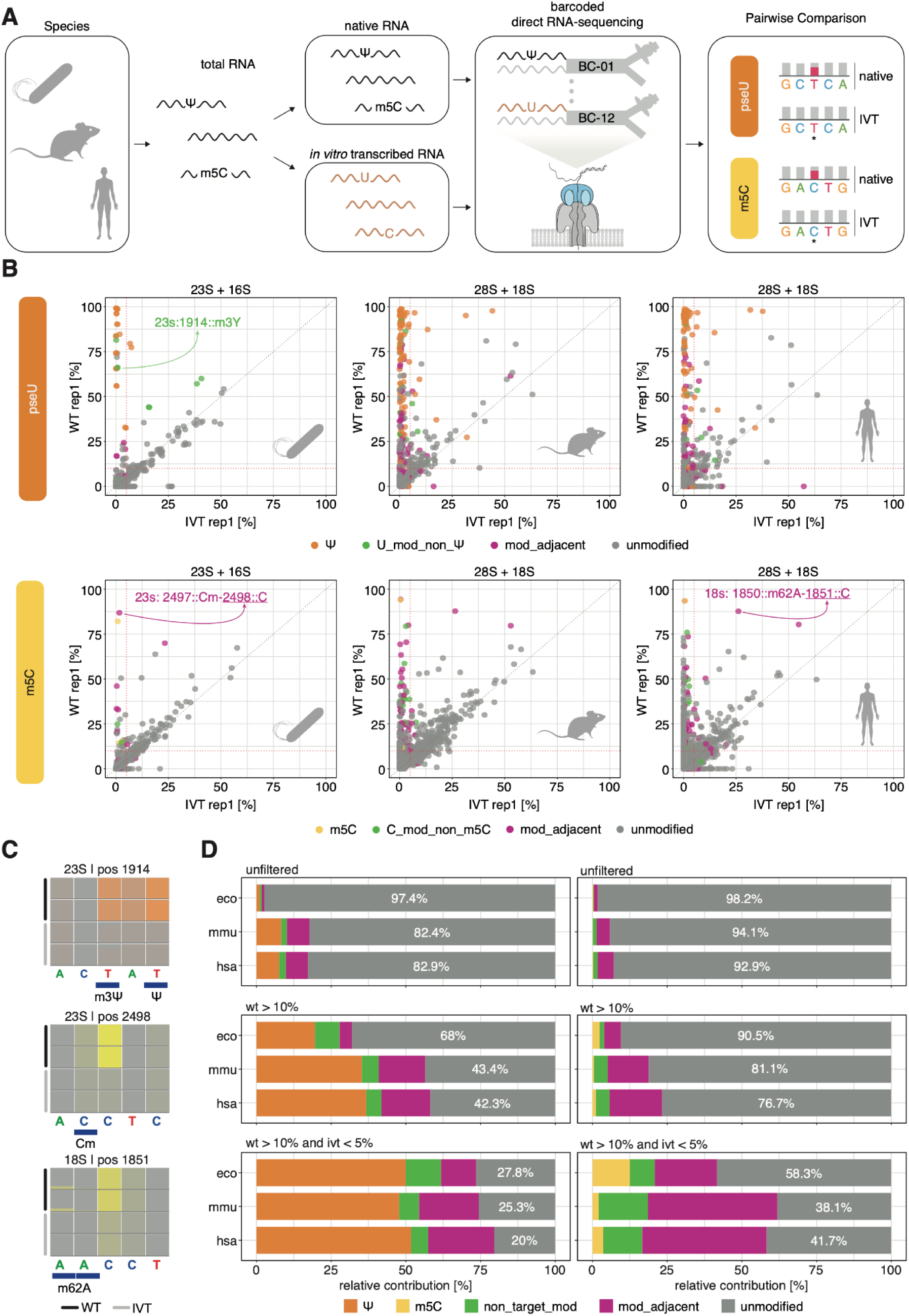
Modification-free controls are essential to remove false positives predicted by ONT modification-aware basecalling models. **(A)** Schematic representation showing the generation of *in vitro* transcribed control samples from the total RNA of three species (*E. coli, M. musculus, and H. sapiens)*, followed by multiplexed sequencing and pairwise comparisons of the basecalled data using either the pseU or m5C model. **(B)** Scatterplots of predicted modification stoichiometry, depicting the relationship of predicted modified sites between the wildtype (WT) and *in vitro* transcribed (IVT) samples for rRNAs found in *E. coli*, *M. musculus,* and *H. sapiens*. Red dotted lines depict the WT and IVT thresholds at > 10% and < 5%, respectively. Individual sites are highlighted to demonstrate the concept of non-targeted modifications affecting the same nucleotide (green) and adjacent modifications found +/−1 nt of the predicted site (purple). **(C)** IGV snapshots of sites highlighted in (B) for both WT and IVT replicates. *Top:* m^3^Ψ mispredicted as pseudouridine (Ψ). *Middle*: canonical C mispredicted as m^5^C due to the presence of C_m_ at position 2497 (23s). *Bottom:* canonical C mispredicted as m^5^C due to the presence of m^6^2A at positions 1849-1850 (18s). Tracks were scaled to a maximum coverage of 500 reads. Each nucleotide position is coloured based on their predicted modification stoichiometry. **(D)** Barplots depicting the relative fractions of true positives and false positives for unfiltered (top), wildtype filtered (middle), and wildtype+ivt filtered (bottom) datasets. False positives have been subdivided into: i) non-target modifications (non_target_mod, green) which correspond to modifications appearing on the same nucleotide as the target modifications. (e.g.: Cm for the m5C model), ii) unmodified sites adjacent to other RNA modification types (e.g., unmodified C neighbouring an m^6,6^A site for the m5C model, shown in purple), and iii) unmodified bases (shown in gray). Percentages of sites predicted as unmodified (gray) are indicated within each bar.

We then specifically assessed whether the use of IVT controls would eliminate all false positives identified in WT sequences. To this end, we compared the predicted stoichiometry of WT and IVT samples basecalled with either the m5C or pseU models (**Fig. 4B**, see also **Tables S5** and **S6**). We found that, while most of the false positive (FP) sites were predicted in both samples – i.e., those found in the diagonal (**Fig. 4B**)–, several FP sites were only detected in the WT sample. Upon closer examination, we observed that some of these FP sites exclusively found in WT samples corresponded to cross-reactivities against modifications affecting the same nucleotide (e.g., m^3^Y in the case of pseU model) (**Fig. 4B-C, and Fig. S6**), in agreement with our previous observations using synthetic RNA molecules (**Fig. 1**). However, we also found that many FP sites corresponded to unmodified residues that had an RNA modification neighboring them (e.g. C_m_ or m^6^ A in the case of the m5C model) (**Fig. 4B-C, and Fig. S6**). This suggests that IVT controls are essential, yet insufficient, to remove FP predictions from basecalling models. Notably, in the case of *H. sapiens* we observed a higher median modification stoichiometry for sites with neighboring modifications than those having a different modified nucleotide from the target one, suggesting that neighboring modifications, even when found at different bases, can be a major confounder when using modification-aware basecallers, causing the neighbouring unmodified base to be predicted as modified (**Fig. S6A**).

Finally, we quantitatively examined how many false positives would be identified in rRNAs after applying a double threshold, requiring a minimum modification frequency on the WT samples (>10%) as well as maximum modification frequency on the IVT samples (<5%). While we observe that the use of these thresholds removes a significant proportion of false positive predictions in both the pseU and m5C models, our results show that the basecalling models still mispredicts as modified ∼48-52% (pseU model) and ∼87-98% (m5C model) of the sites (**Fig. 4D**) following the double threshold filtering step. Altogether, our work shows that the use of IVT samples allows significant removal of false positives; however, it also identifies a previously disregarded source of false positives that can neither be eliminated by the inclusion of additional modifications in the model, nor by the use of IVT controls (**Fig. 4D**).

### Identification of modifications in the absence of modification-aware basecalling models

While modification-aware basecalling models open the possibility to identify modifications in individual molecules, this approach is only applicable to a handful of RNA modifications for which a model is currently available. Therefore, most RNA modifications described to date (43) cannot be identified using modification-aware basecalling models, leaving the majority of RNA modification types unidentified in DRS datasets. To address this, here we examined whether previous approaches, such as the use of ‘systematic basecalling errors’ (8, 10, 12, 16, 23), and altered current intensity features (e.g. signal intensity and dwell time) (13, 16, 28) can be employed to identify RNA modifications using the RNA004 chemistry.

Previous works have shown that basecalling ‘error’ patterns are significantly affected by the choice of canonical model used to basecall the reads (21, 27). Therefore, we first examined whether the different canonical models released for dorado (*fast*, *hac* and *sup*) would capture different RNA modification types, using synthetic ‘curlcakes’ sequenced with RNA004 chemistry (**Fig. 5A, see also Table S7**). Our results demonstrate that the *fast* model generates highest global ‘error’ rates, compared to *hac* and *sup* models at unmodified sites (**Fig. 5B**). Moreover, all RNA modifications could be detected in the form of basecalling ‘errors’ across all models. However, these ‘errors’ were significantly decreased in the case of Ψ, m^5^C and m^6^A modifications both in *hac* and *sup* models, with their error rate being close to the background error rate of the unmodified bases (**Fig. 5B**), suggesting that these modifications have likely been included in the training of ‘canonical’ models (i.e., the canonical *hac* and *sup* models have learnt to basecall ‘m^6^A’ as ‘A’). By contrast, ac^4^C, m^1^Ψ and m^5^U modifications produced high basecalling ‘errors’ compared to their unmodified counterparts (ΔSumErr), confirming that base-calling ‘errors’ can be used to detect RNA modifications for which no modification-aware basecalling model exists (**Fig. 5C**).

**Figure 5.**
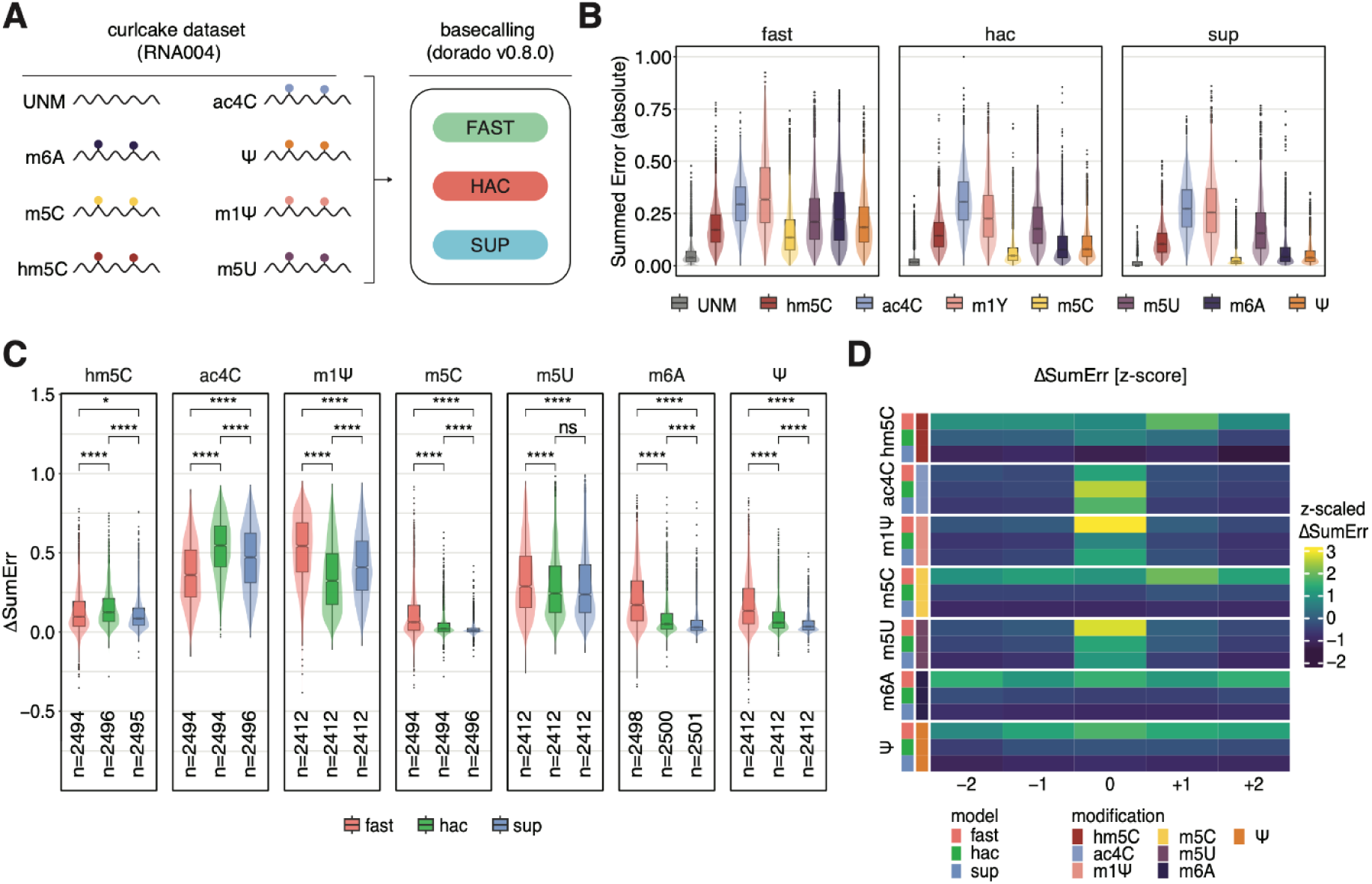
Prediction of RNA modifications using systematic basecalling errors in RNA004 chemistry. **(A)** Schematic representation of the ‘curlcake’ molecules used to benchmark the use of basecalling ‘errors’ to detect RNA modifications, using either fast, hac or sup canonical dorado models. **(B)** Summed Errors of unmodified and modified bases of curlcake datasets basecalled with either *fast* (left), *hac* (middle) or *sup* (right) dorado canonical basecalling models. **(C)** Comparative analysis of the differential summed error (ΔSumErr) for each RNA modification type examined, calculated by subtracting the ‘error’ signal obtained at modified and unmodified sites, for each possible 5-mer sequence. **(D)** Analysis of the position at which the ‘error’ signature is found, for each RNA modification type (Ψ, m^6^A, m^5^U, m^5^C, m^1^Ψ, ac^4^C and hm^5^C) and canonical basecalling model (*fast*, *hac*, *sup*).

Next, we examined whether the ‘error’ pattern would be centered in the modified site, or in a neighbouring position. To this end, we aggregated the ΔSumErr from all 5-mers, and examined their ‘error’ patterns along each position of the 5-mer (with 0 corresponding to the modified site) (**Fig. 5D**). Our results show that, globally, most RNA modifications showed ‘error’ patterns at the modified site (pos 0), whereas m^5^C and hm^5^C showed increased errors at the neighbouring position (+1), in agreement with what had been previously observed with RNA002 chemistry (16, 27). While the fast model produced the highest ‘error signal (ΔSumErr) for most RNA modification types, we should note that this was not the case for ac^4^C modifications (**Fig. 5B-D**), suggesting that if future users are interested in identifying this modification type, *hac* models will likely better capture their signal.

Finally, we sequenced total RNA from *E. coli* wild type and knockout strains (KO) lacking specific rRNA modifications at known positions: rsmA KO (m^6,6^A at 16S:1518 and1519), rsmG KO (m^7^G at 16S:527), rsmD KO (m^2^G at 16S:966) and rsmE (m^3^U at 16S:1498) (see *Methods*) (**Fig. 6A**, see also **Table S8**). We then examined how these RNA modifications would be detected in the form of ‘errors’ using by comparing the basecalling ‘error’ patterns of wild type and knockout samples, as well as in the form of altered current features (signal intensity and dwell time) (see *Methods*). Our results showed that all four modification types examined could be robustly identified in the form of basecalling errors (ΔSummedError), alterations in current intensity (ΔSignal Intensity) as well as alterations in dwell time (ΔDwell Time) (**Fig. 6B)**. Thus, we conclude that classical approaches relying on pairwise comparisons remain an effective approach to identify differential RNA modifications by direct comparison of the features from two samples, in agreement with previous observations obtained using RNA002 chemistry (44).

**Figure 6.**
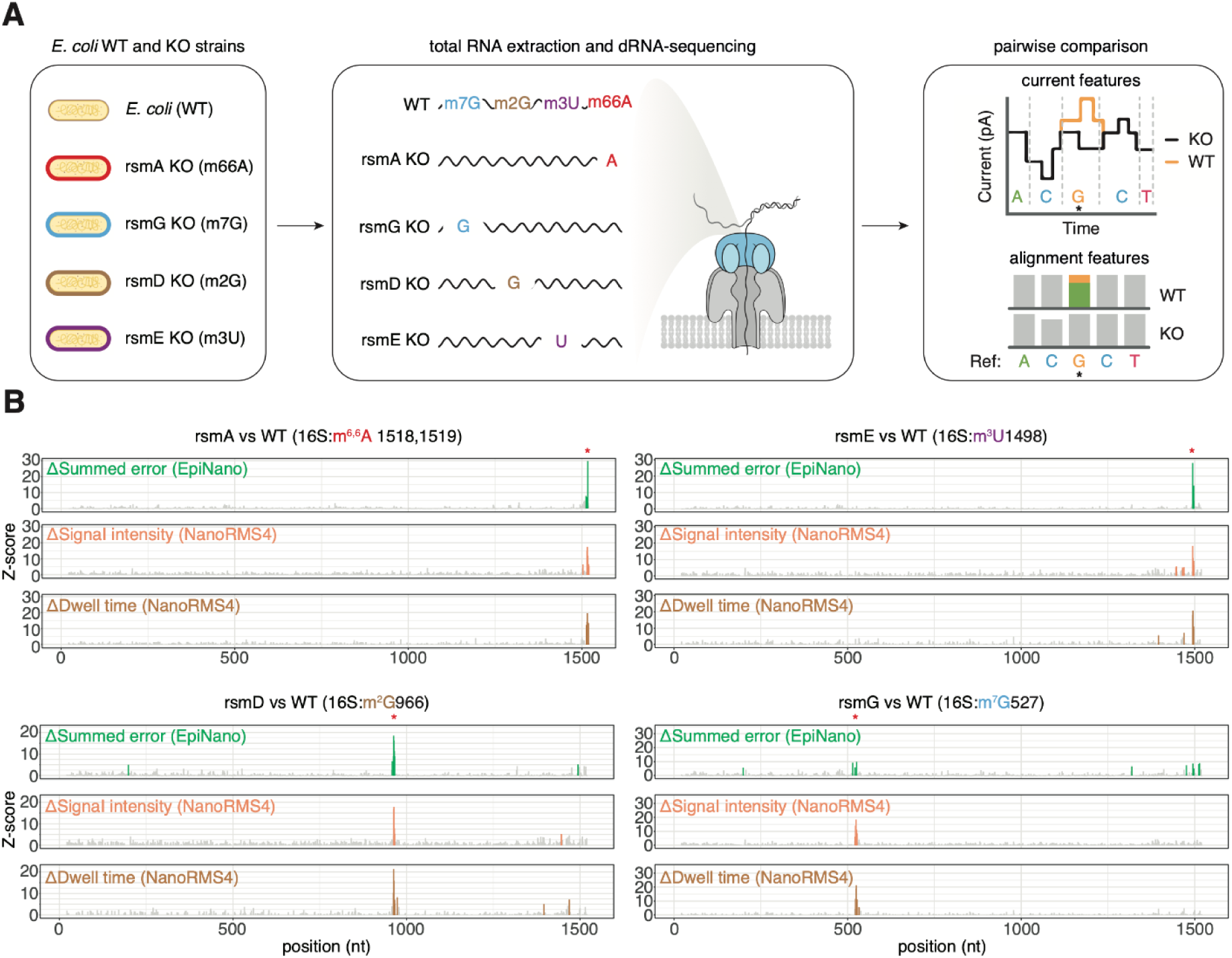
Prediction of RNA modifications using differential basecalling errors and alterations in raw nanopore current features in RNA004 chemistry. **(A)** Schematic representation depicting the workflow used to analyse *E.coli* wild type and knockout strains. Modifications placed by the indicated enzymes, for which not modification-aware basecallers exist to-date, are shown in brackets. **(B)** Per-position differential modification Z-scores along the 16S rRNA transcript, comparing the modification profiles of E. coli wild type and each knockout (KO) strain. For each pairwise comparison, the differential modification profiles were obtained by computing: i) ΔSummed Error (green), computed using *EpiNano*; ii) ΔSignal intensity (orange), using *NanoRMS4*, and iii) ΔDwell Time (brown), using NanoRMS4. The expected differentially modified site for each pairwise comparison is highlighted with a red asterisk.

## DISCUSSION

The emergence of native RNA sequencing has enabled the possibility to examine diverse RNA modifications simultaneously, in full-length reads, with single nucleotide and single molecule resolution. This has been greatly advanced by the appearance of modification-aware basecalling models, which allow the detection of RNA modifications during the basecalling step. Following initial attempts by the scientific community (18, 21, 29, 45), ONT recently released several modification-aware modifications for a handful of RNA modification types. However, their performance, cross-reactivities and false positive rates have only recently started to be explored (46–50).

Here, we systematically examined the performance of modification-aware ONT basecalling models using a panel of synthetic constructs covering all possible 5-mers, and containing diverse RNA modification types, both cognate and non-cognate to the models in question, as well as on *in vivo* and IVT samples from *E. coli,* S*. cerevisiae, M. musculus* and *H.sapiens*. Our results demonstrate that base-calling models capture most annotated modified sites; however, they mispredict a significant number of sites as modified, leading to an overall high proportion of false positives (**Fig. 3**). While we demonstrate that IVT controls can significantly reduce the number of false positives (**Fig. 4**), we show that a large proportion of incorrectly predicted sites still remain after a double filtering step (**Fig. 4**). These potential mispredictions could lead to incorrect conclusions when interpreting the biological results, and could further increase controversies in the RNA modifications field, specially in scenarios in which the existence of RNA modifications in certain RNA biotypes has been questioned (42, 51, 52). Therefore, caution must be taken when interpreting the results to acquire novel biological insights of previously uncharacterized RNA species.

There is a growing consensus about the need to incorporate modification-free transcriptome-wide in vitro controls (IVT) as negative controls to improve the accuracy of RNA modification predictions (21, 22, 40). This is true not only for nanopore-based approaches (12, 42, 53), but also for methods coupled to next-generation-sequencing (40, 54). Incorporating such controls will likely become routine practice in most laboratories doing RNA modification analyses. We should note however, that in our work we identify three main sources of false positives by modification-aware basecalling models, some of which cannot be removed using IVT controls: i) those inherent to the k-mer context, which can largely be removed using IVT controls (**Fig. 4B**); ii) those caused by cross-reactivity with other RNA modification types that exist at the same reference base, which cannot be removed with IVT controls –but could be potentially removed in the future once modification-aware basecalling models for additional modifications become available– (**Fig. 1, 4B-C**); and iii) those caused by cross-reactivity with other RNA modification types present at neighbouring positions, causing misprediction of a modified residue at an unmodified site (e.g., m^6,6^A neighboring an unmodified C, leading to m^5^C misprediction by the m5C model (**Fig. 4C**). Notably, this latter type of false positives will remain even if modification-aware models incorporate additional modifications. This can significantly limit the applicability of modification-aware base-calling models in highly modified RNA biotypes, such as tRNA, rRNA or synthetic RNA constructs.

A key strength of the RNA004 chemistry, in addition to the availability of ONT-supported RNA modification-aware base-calling models, is a major increase in sequencing yield, compared to the previous RNA002 chemistry. Consequently, there is a pressing need to multiplex DRS runs. While ONT recently announced the availability of DRS multiplexing kits in the near future, to date this is not yet available, and the number of barcodes to be released remains unclear. Here, we provide a pre-trained 96-barcode demultiplexing model compatible with the latest RNA004 chemistry, enabling high-multiplexing of DRS RNA004 sequencing runs. The possibility to multiplex up to 96 barcodes in a single sequencing flowcell does not only improve the cost-effectiveness of DRS sequencing, but also significantly reduces batch effects across flowcells and samples, as these are otherwise sequenced and prepared independently. Moreover, the possibility of multiplexing enhances the incorporation of control samples by future users, such as IVT conditions, ensuring that future works using modification-aware basecalling models can also reduce false positives in their future experiments.

## METHODS

### *E. coli* culturing and total RNA extraction

*E. coli* strains used in this work were obtained from the Keio Knockout Collection (55), including the reference wild type strain (BW25113) (Cultek). Knockout strains have a kanamycin cassette replacing the depleted gene. Strains were plated in LB-agar plates (*E. coli* BW25113) or in LB-agar plates supplemented with 25 μg/mL kanamycin (in the case of *E. coli* knockout strains). For each strain, starter cultures were grown in 4 mL LB, incubated at 37 °C and 180 rpm. After 6 h, 4 ml cultures were centrifuged in a pre-chilled benchtop centrifuge at 16,000 g 4 °C for 5 min. Supernatant was discarded, and pellets were stored at −80 °C overnight. Frozen pellets were thawed on ice, followed by the addition of 400 μL of Trizol (Life Technologies, #15596018). After 5 minutes incubation at room temperature, 90 μL of chloroform (Vidra Foc, #C2432) was added and mixed thoroughly by inversion. Samples were centrifuged for 15 min at 16,000 g at 4 °C. Supernatant was kept and an equal volume of 70% ethanol was added. Afterwards, small RNAs were removed with the Qiagen RNeasy Mini Kit following manufacturer’s recommendations (Qiagen, #74104). Samples were DNAse-treated using Turbo DNAse (Life Technologies, #AM2239) for 15 min at 37 °C. Finally, the sample was cleaned with 1X AMPure RNAClean XP beads (Beckman Coulter, A63987). RNA integrity was assessed using TapeStation and quantified using Nanodrop.

### *S. cerevisiae* culturing and total RNA extraction

*S. cerevisiae* strains (BY4741) were grown in 4 ml of standard YPD medium (1% yeast extract, 2% Bacto Peptone and 2% dextrose) overnight at 30 °C. The next day, cultures were diluted to 0.0001 OD_600_ in 200 ml of YPD and grown overnight at 30 °C with shaking (250 r.p.m.). When cultures reached the mid-exponential growth phase (OD_600_ 0.5), the cultures were transferred into a pre-chilled 50-ml Falcon tube and centrifuged at 3,000g for 5 minutes at 4 °C, followed by two washes with water, and then pellets were snap-frozen at −80 °C. Snap-frozen yeast pellets were resuspended in 660 µl of TRIzol Reagent (Thermo Fisher Scientific, 15596018) with 340 µl of acid-washed and autoclaved 425–600-µm glass beads (Sigma-Aldrich, G8772). The cells were disrupted by vortexing on top speed for seven cycles of 15 seconds and chilling the samples on ice for 30 seconds between cycles. The samples were then incubated at room temperature for 5 minutes, and 200 µl of chloroform was added. After briefly vortexing the suspension, the samples were incubated for 5 minutes at room temperature and centrifuged at 14,000g for 15 minutes at 4 °C. The upper aqueous phase was transferred to a new tube. To precipitate RNA, 1× volume of molecular-grade isopropanol and 1 µl of GlycoBlue co-precipitant (Thermo Fisher Scientific, AM9515) were added and mixed by inverting and incubated for 10 minutes at room temperature. The samples were centrifuged at 14,000g for 15 minutes at 4 °C, and the pellet was then washed with ice-cold 70% ethanol. The pellet was resuspended in 20 µl nuclease-free water after air drying for 5 minutes on the benchtop, and the RNA purity was measured using a NanoDrop 1000 spectrophotometer. Then, 10 µg of each sample was treated with Turbo DNase (Thermo Fisher Scientific, AM2238) and subsequently cleaned up using a Zymo RNA Clean and Concentrator-5 kit (Zymo Research, R1016) following the manufacturers’ instructions The RNA was eluted in nuclease-free water. RNA concentration was determined using Qubit Fluorometric Quantitation; RNA purity was measured with a NanoDrop 1000 spectrophotometer; and the RNA electropherogram was obtained using Agilent 4200 TapeStation RNA HS ScreenTape Assay.

### Mouse models and total RNA extraction

The mice used in this study were of the C57BL/6J strain (Charles River, strain #000664). Animals were maintained on a defined control diet (Special Diets Services, RM1 (P), 801151), housed in standard cages at 22–24°C, and given ad libitum access to food and water. Lighting conditions were regulated using a 12:12 hour light-dark cycle, with lights turned off at 19:00. Adult C57BL/6J mice were euthanized using CO₂. All tissues were promptly dissected, snap-frozen in liquid nitrogen, and stored at −80°C until further use. Animal experiments were conducted in accordance with EU Directive 86/609/EEC and Recommendation 2007/526/EC concerning the protection of animals used for scientific purposes, as implemented by Spanish law 1201/2005. All procedures were approved by the institutional ethics committee.

### Human cell line culturing conditions and total RNA extraction

MCF7 (ATCC) cell lines were cultured in Gibco Life Technologies Dulbecco’s modified Eagle medium (DMEM), supplemented with 10% (v/v) fetal bovine serum (FBS), and 1% Penicillin Streptomycin. In order to extract RNA from MCF7 cell cultures, they were grown in 9.6 cm2 plates at 37°C in a humidified atmosphere containing 5% CO2 until. Once cells reached 80% confluency, they were first washed with PBS once and then incubated with 1 ml Trizol at room temperature for 5 min. For each ml of trizol, 200 μl of chloroform were added, sampled were vortexed for 10 s, incubated for 2-3 min at RT, and spun down at 16,000 x g for 15 min at 4°C. The clear top phase was transferred to a new tube, and purified with the miRNeasy kit (Qiagen) according to the manufacturer’s instructions. The eluted RNA was quantified by Qubit and RNA integrity was assessed by Agilent Tapestation.

### Generation of *in vitro* transcribed rRNAs from total RNA

We generated *in vitro* transcribed rRNA following previously published literature with some minor adjustments (12, 22). First, total RNA was pre-fragmented to ensure equal coverage across long transcripts by using 150 ng of total RNA, 2µl of RNA Fragmentation Buffer (NEB, E6150S) in a total volume of 20µl. The reaction was incubated for 1min at 94°C to obtain a fragment size of ∼500nt, and stopped by the addition of 2µl 10X RNA Fragmentation Stop Solution (NEB, E6150S) and transfer to 4°C. Next, 3’ ends were prepared by the addition of 4µl of FastAP (Thermo Scientific, EF0651), 8µl of 10X Fast-AP Buffer, in a total volume of 80µl. The reaction was incubated at 37°C for 15min followed by a cleanup using the Zymo Clean and Concentrator-5 kit (Zymo Research, R1016). Finally, a short *in-vitro* polyadenylation combining the pre-fragmented total RNA with 1µl of *E.coli* Poly(A) Polymerase (NEB, M0276S), 2µl of ATP (10mM), and 2µl of 10X Poly(A) Polymerase Buffer, in a total of 20µl was carried out. The reaction was incubated at 37°C for 2min followed by a cleanup using the Zymo Clean and Concentrator-5 kit.

For the reverse transcription reaction, 50ng of input RNA was combined with 1µl of dNTPs (NEB, N0447S), and 1µl of the CRTA primer (2µM, 5’-ACTTGCCTGTCGCTCTATCTTCTTTTTTTTTT-3’) in a total volume of 11µl, followed by an incubation at 65C for 5min before being snap cooled to 4C. Next, 4µl of 5X RT Buffer (Thermo Scientific, EP0752), 1µl of RNAseOUT (Thermo Scientific, 10777019), 1µl of ddH20, and 2µl of the SSPII primer (10µM, 5’-TTTCTGTTGGTGCTGATATTGCTTTVVVVTTVVVVTTVVVVTTVVVVTTTmGmGmG-3’) were added. The reaction was incubated at 42°C for 2min before 1µl Maxima H Minus RT (Thermo Scientific, EP0752) was added. The reverse transcription reaction was incubated at 42°C for 90min followed by heat inactivation at 85°C for 5min. Next, a PCR reaction was performed to enrich for transcripts that underwent successful reverse transcription and template switching. To this end, 5µl of cDNA from the previous step was added to 25µl of LongAmp® Taq 2X Master Mix (NEB, M0287S), 2µl of T7_fwd primer (10µM, 5’-TAATACGACTCACTATAGCGAGGCGGTTTTCTGTTGGTGCTGATATTGCT-3’), and 2µl of T7_rev primer (5’-ACTTGCCTGTCGCTCTATCTTC-3’) in a total reaction volume of 50µl. The PCR consisted of a initial denaturation step at 95°C for 5min followed by 12 cycles of denaturation (95°C, 15s), annealing (60°C, 15s) and extension (65°C, 15min), with a final extension step at 65°C for 15min. To remove excess T7_fwd primer, 1µl of Exonuclease was added and the reaction incubated at 37°C for 15min, followed by heat inactivation at 80°C for 15min. Amplified cDNA was cleaned up using 1.8X AMPure XP beads (Beckman Coulter, A63880). Eluted cDNA was used to set up an overnight (16h) *in vitro* transcription using the AmpliScribe T7 High Yield Transcription Kit (Biosearch Technologies, AS3107), following the manufacturer’s instructions. IVT libraries were cleaned up using the RNEasy Mini Kit (Qiagen, 74104) following the manufacturer’s instructions. To make IVT libraries amenable to nanopore sequencing library preparation, a short *in vitro* polyadenylation was performed using 5µg of IVT library, 1µl of *E.coli* Poly(A) Polymerase (NEB, M0276S), 2µl of ATP (10mM), and 2µl of 10X Poly(A) Polymerase Buffer, in a total volume of 20µl followed by cleanup using the Zymo Clean and Concentrator-5 kit.

### Generation of synthetic RNA molecules containing all possible 5-mer sequences

This study used synthetic RNAs known as ‘curlcakes’, which were designed to include all possible 5-mer sequences while minimizing RNA secondary structures (7), and consist of four *in-vitro* transcribed constructs: (1) Curlcake 1, 2,244 bp; (2) Curlcake 2, 2,459 bp; (3) Curlcake 3, 2,595 bp and (4) Curlcake 4, 2,709 bp. To generate ‘curlcake’ synthetic RNAs, plasmids (pUC57) were digested with BamHI-HF (NEB, R3136L) and EcoRV-HF (NEB, R3195L) as per manufacturer instructions, followed by a clean-up using phenol/chloroform/isoamyl-alcohol 25/24/1, v/v, pH = 8.05 (Sigma Aldrich, P3803). Linearized plasmids were used for overnight *in vitro* transcription with the AmpliScribe T7-Flash Transcription Kit (Lucigen, ASF3507). To generate fully modified ‘curlcakes’, canonical nucleotides were exchanged for the following modified nucleotides: pseudouridine-5’-Triphosphate (pseudo-UTP, Trilink, N-1019) instead of UTP, 5-Methyluridine-5’-Triphosphate (5-meUTP, Trilink, N-1024-1) instead of UTP, *N1*-Methylpseudouridine-5’-Triphosphate (me1Ψ-UTP, Jena Bioscience, NU-890S) instead of UTP, 5-methylcytidine-5’-Triphosphate (5-methyl-CTP Trilink, N-1014), 5-Hydroxymethylcytidine-5’-Triphosphate (5-hme-CTP, Trilink, N-1087), *N4*-Acetyl-cytidine-5’-Triphosphate (*N4*-Acetyl-CTP, Jena Bioscience, NU-988S) instead of CTP or *N6*-methyladenosine-5’-Triphosphatetriphosphate (*N6*-Methyl-ATP, Trilink, N-1013). To generate ‘curlcakes’ containing different overall modification levels, mixtures of canonical and modified NTPs were used during *in vitro* transcription (e.g., 1/4 *N6*-Methyl-ATP, 3/4 ATP, to obtain 25% m^6^A residues). Following *in-vitro* transcription, products were treated with Turbo DNase (Thermo Fisher, AM2238) to remove the template plasmid and cleaned up using the RNeasy MinElute Kit (Qiagen, 74204).

### Nanopore direct RNA sequencing of IVTs and total RNA samples

IVT constructs and total RNA samples were polyadenylated using Escherichia coli poly(A) polymerase (NEB, M0276S) according to the manufacturer’s instructions followed by a clean-up using the RNA Clean & Concentrator-5 kit (Zymo Research, R1013). Concentrations were determined using the Qubit 2.0 Fluorometer. RNA integrity, and polyadenylation success, were determined using an Agilent 4200 TapeStation. Library preparation was carried out according to the manufacturer’s instructions (direct-rna-sequencing-sqk-rna004-DRS_9195_v4_revB_20Sep2023-promethion) with minor adjustments to enable highly multiplexed sequencing of multiple barcoded samples (25). Adapter ligation was carried out for 15 min at room temperature using 100 ng of poly(A)-tailed substrate, 0.5 µl of custom barcode adapter (1.4 µM), 2 µl of NEBNext® Quick Ligation Reaction Buffer (NEB, B6058S), 0.5 µl of SUPERase·In RNAse Inhibitor (Invitrogen, AM2696) and 0.5 µl of concentrated T4 DNA Ligase (NEB, M0202T) in a total volume of 10 µl nuclease-free H_2_O. Subsequently, 4 µl of 5×SuperScript IV Buffer, 1µl of dNTPs (NEB, N0447S), 2 µl 0.1 DTT, and 1 µl of SuperScript™ IV Reverse Transcriptase (Thermo Fisher, 18090010) were added and the reaction incubated for 15 min at 50°C before heat inactivation for 5 min at 70°C. Then, RNA:DNA samples designated for sequencing in the same run were pooled accordingly. Separate pools were prepared based on the number of sequencing runs required, and each pool was cleaned up using 1x RNAClean XP beads (Beckman Coulter, A63987) and eluted in nuclease-free H_2_O. 23 ul of eluate per pool (aprox. 1 µg) was ligated to 6 µl RNA Ligation Adapter (RLA) with 3 µl T4 DNA Ligase (NEB, M0202T) and 8 µl of NEBNext® Quick Ligation Reaction Buffer (NEB, B6058S) in a total volume of 40 µl, by incubating at room temperature for 15 min. Libraries were cleaned up using 0.6x RNAClean XP beads (Beckman Coulter, A63987), washed twice with 150 µl of Wash Buffer (WSB) and eluted in 33 µl of elution buffer (EB). 1 µl of each library was used for quantification with Qubit™ 1X dsDNA High Sensitivity (HS) (Invitrogen, Q33231) and 32 µl were mixed with 100 µl of Sequencing Buffer (SB) and 68 µl of Library Solution (LIS). Libraries were run on a primed FLO-PRO004RA flowcell using a PromethION 2 solo sequencer with MinKNOW acquisition software version v24.11.10.

### Preprocessing of raw nanopore direct RNA sequencing data

Preprocessing of raw nanopore sequencing data (pod5) was performed using the Master of Pores nextflow pipeline version 4 (MoP4) (56). First, as part of the mop_preprocess workflow, the sequencing data was first demultiplexed with SeqTagger (v.1.0c) using the 96 barcode model developed for RNA004 (`-k b96_RNA004`) using the default baseQ cutoff of > 50 (`-b 50`) (25). Next, reads were basecalled using *dorado* (v0.8.0) using one of the modification-aware basecalling models (m6A_DRACH, inosine_m6A, pseU and m5C) with either the hac or sup accuracy settings (v5.1.0). Basecalled reads were aligned to their respective reference using minimap2 (v2.17) (57) with default settings (`-y -ax map-ont -k 5`). Finally, QC plots were generated using NanoPlot (v1.32.1) (58) to visualise read-length vs. average-read quality, and custom R scripts (R version 4.3.2) for coverage plots (**Fig. S2**, *see also* **Table S2**). The b96_RNA004 model is openly available on GitHub (https://github.com/novoalab/SeqTagger).

### Per read modification probability analysis on synthetic constructs

Per-read and position modification probabilities were extracted for each alignment file (BAM) generated by the modification-aware basecalling models using modkit (v0.4.2, https://github.com/nanoporetech/modkit). For the analysis of the *in vitro* data, 10,000 reads were subsampled from each alignment file (`modkit extract full --mapped-only --num-reads 10000 <input.bam> <output.tsv>`). For the analysis of the in vivo data, the following command was used: `modkit pileup “$BAM_FILE” “$BED_OUTPUT” --motif C 0 --ref “$REFERENCE” --log-filepath “$LOG_FILE”` (motif C was used for the m5C basecaller and motif T for the Ψ basecaller). The resulting tables were further processed using custom R scripts to generate modification probability plots, ROC and PR-ROC plots (**Fig. 1**). Reads with a minimum baseQ of < 10 were excluded from the analysis. To perform motif enrichment analysis, we used STREME (v5.5.7) (59) with the entire set of kmers present in curlcakes as the background model and and the underestimated set as test set (modification probability < 0.75, **Fig. 2B**) (`streme --verbosity 1 --oc. --rna --totallength 4000000 --time 14400 --minw 5 --maxw 5 --thresh 0.05 --align center`).

### Per site stoichiometric analysis of synthetic constructs

To perform per-site analysis, `modkit pileup` was run with the default filter-threshold as part of the mop_mod MasterOfPores4 workflow (`modkit pileup --with-header --filter-percentile 0.1`). For the m6A_DRACH model, pileup was run with the additional motif flag (-`-motif DRACH 2 --ref <ref.fasta>`) to ensure that only predictions for the mammalian m^6^A consensus sequence are considered. Next, predictions were filtered for the plus strand, and results stemming from different barcodes were collapsed into a single master table using bedtools unionbedg (v2.31). Finally, stoichiometric analysis across a range of modified fractions was performed using custom R scripts (**Fig. 2C**).

### Per site stoichiometric analysis of native and *in vitro* transcribed rRNA

To perform per-site analysis, `modkit pileup` was run with the default filter-threshold (`modkit pileup --with-header --filter-percentile 0.1 --motif N 0`, where N represents the target base). Finally, modkit tables were filtered using custom R scripts to filter for sites with a minimum valid coverage (N_val_) of > 500 reads. Tables were further annotated with known modified sites. Annotations can be found on the project’s github page (https://github.com/novoalab/ONT_basecalling_models/tree/main/ref/annotations). Species silhouettes were obtained from PhyloPics (https://www.phylopic.org/). All scripts have been deposited on the project’s github page (https://github.com/novoalab/ONT_basecalling_models/tree/main/).

### Alignment and current feature analysis of synthetic constructs and E.coli rRNA sequences

To identify RNA modifications in the form of systematic basecalling ‘errors’, POD5 files from synthetic ‘curlcake’ datasets and from total *E. coli* RNA samples were processed using the MasterOfPores4 workflow, using either *fast*, *hac* or *sup* dorado (https://github.com/nanoporetech/dorado) canonical models (v0.8). Modifications were predicted using the EpiNano software (23) to extract alignment features or NanoRMS4 (16) to extract current features (delta signal intensity and dwell time) using the mop_mod module. Differential basecalling errors were converted into Z-scores using the mop_consensus module (44).

## ANNOTATIONS

The list of rRNA modifications present in *E. coli*, *S. cerevisiae, M. musculus,* and *H. sapiens* rRNAs was obtained from MODOMICS (https://iimcb.genesilico.pl/modomics/sequences/) (43). Annotations and reference sequences for all species used in this work were taken from are available on GitHub (https://github.com/novoalab/ONT_basecalling_models/tree/main/ref).

## DATA AVAILABILITY

Data supporting the findings of this study are available from the corresponding author (EMN), upon reasonable request. (see also **Table S2**).

## CODE AVAILABILITY

Code to analyze modification-aware basecalled data has been deposited in GitHub (https://github.com/novoalab/ONT_basecalling_models/tree/main).

## AUTHOR CONTRIBUTIONS

LLL generated the *in vitro* ‘curlcake constructs’, cultured *E. coli* wild type and knockout strains and extracted the total RNA. GD generated training data for the 96-barcode model and *in vitro* libraries from total RNA. LLL and GD prepared and sequenced the samples. GD performed the bioinformatic analysis of *in vitro* curlcake constructs. IM performed the bioinformatic analysis of the in vivo samples (*E. coli* and *S. cerevisiae*). GD and IM performed comparative analysis of wildtype and *in vitro* transcribed rRNA. FP and AM analyzed the synthetic ‘curlcake’ and in vivo datasets using base-calling ‘errors’ on the canonical modification-unaware models. GD, IM and EMN built the figures. EMN conceived and supervised the project. GD, IM, and EMN wrote the manuscript, with contributions from all authors.

## ACKNOWLEDGEMENTS

We thank all the members of the Novoa lab for their insightful discussions. This project has received funding from the European Union’s Horizon 2020 research and innovation programme under the Marie Sklodowska-Curie grant agreement No 956810 (GD). EMN and this project received funding from the project PID2021-128193NB-I00 through the MCIN/AEI/10.13039/501100011033/ FEDER, UE; the European Union’s Horizon Europe through the European Research Council under the grant agreement number 101042103 and 101187456. Views and opinions expressed are however those of the author(s) only and do not necessarily reflect those of the European Union. Neither the European Union nor the granting authority can be held responsible for them. We acknowledge support of the Spanish Ministry of Science and Innovation through the Centro de Excelencia Severo Ochoa (CEX2020-001049-S, MCIN/AEI /10.13039/501100011033), and the Generalitat de Catalunya through the CERCA programme and to the EMBL partnership. We are grateful to the CRG Core Technologies Programme for their support and assistance in this work.

## COMPETING INTERESTS

EMN is a member of the Scientific Advisory Board of IMMAGINA Biotech. GD, IM and EMN have received travel bursaries from ONT to present their work at conferences.

## REFERENCES

1. Frye, M., Jaffrey, S.R., Pan, T., Rechavi, G. and Suzuki, T. (2016) RNA modifications: what have we learned and where are we headed? Nat. Rev. Genet., 17, 365–372.

2. Suzuki, T. (2021) The expanding world of tRNA modifications and their disease relevance. Nat. Rev. Mol. Cell Biol., 22, 375–392.

3. Garalde, D.R., Snell, E.A., Jachimowicz, D., Sipos, B., Lloyd, J.H., Bruce, M., Pantic, N., Admassu, T., James, P., Warland, A., et al. (2018) Highly parallel direct RNA sequencing on an array of nanopores. Nat. Methods, 15, 201–206.

4. Diensthuber, G. and Novoa, E.M. (2025) Charting the epitranscriptomic landscape across RNA biotypes using native RNA nanopore sequencing. Mol. Cell, 85, 276–289.

5. Fleming, A.M., Bommisetti, P., Xiao, S., Bandarian, V. and Burrows, C.J. (2023) Direct nanopore sequencing for the 17 RNA modification types in 36 locations in the E. coli ribosome enables monitoring of stress-dependent changes. ACS Chem. Biol., 18, 2211–2223.

6. Jain, M., Olsen, H.E., Paten, B. and Akeson, M. (2016) The Oxford Nanopore MinION: delivery of nanopore sequencing to the genomics community. Genome Biol., 17, 239.

7. Liu, H., Begik, O., Lucas, M.C., Ramirez, J.M., Mason, C.E., Wiener, D., Schwartz, S., Mattick, J.S., Smith, M.A. and Novoa, E.M. (2019) Accurate detection of m6A RNA modifications in native RNA sequences. Nature Communications, 10, 1–9.

8. Abebe, J.S., Price, A.M., Hayer, K.E., Mohr, I., Weitzman, M.D., Wilson, A.C. and Depledge, D.P. (2022) DRUMMER—rapid detection of RNA modifications through comparative nanopore sequencing. Bioinformatics, 38, 3113–3115.

9. Naarmann-de Vries, I.S., Zorbas, C., Lemsara, A., Piechotta, M., Ernst, F.G.M., Wacheul, L., Lafontaine, D.L.J. and Dieterich, C. (2023) Comprehensive identification of diverse ribosomal RNA modifications by targeted nanopore direct RNA sequencing and JACUSA2. RNA Biol., 20, 652–665.

10. Jenjaroenpun, P., Wongsurawat, T., Wadley, T.D., Wassenaar, T.M., Liu, J., Dai, Q., Wanchai, V., Akel, N.S., Jamshidi-Parsian, A., Franco, A.T., et al. (2021) Decoding the epitranscriptional landscape from native RNA sequences. Nucleic Acids Res., 49, e7.

11. Parker, M.T., Knop, K., Sherwood, A.V., Schurch, N.J., Mackinnon, K., Gould, P.D., Hall, A.J., Barton, G.J. and Simpson, G.G. (2020) Nanopore direct RNA sequencing maps the complexity of Arabidopsis mRNA processing and m6A modification. Elife, 9.

12. Tavakoli, S., Nabizadeh, M., Makhamreh, A., Gamper, H., McCormick, C.A., Rezapour, N.K., Hou, Y.-M., Wanunu, M. and Rouhanifard, S.H. (2023) Semi-quantitative detection of pseudouridine modifications and type I/II hypermodifications in human mRNAs using direct long-read sequencing. Nat. Commun., 14, 334.

13. Leger, A., Amaral, P.P., Pandolfini, L., Capitanchik, C., Capraro, F., Miano, V., Migliori, V., Toolan-Kerr, P., Sideri, T., Enright, A.J., et al. (2021) RNA modifications detection by comparative Nanopore direct RNA sequencing. Nat. Commun., 12, 7198.

14. Pratanwanich, P.N., Yao, F., Chen, Y., Koh, C.W.Q., Wan, Y.K., Hendra, C., Poon, P., Goh, Y.T., Yap, P.M.L., Chooi, J.Y., et al. (2021) Identification of differential RNA modifications from nanopore direct RNA sequencing with xPore. Nat. Biotechnol., 39, 1394–1402.

15. Hendra, C., Pratanwanich, P.N., Wan, Y.K., Goh, W.S.S., Thiery, A. and Göke, J. (2022) Detection of m6A from direct RNA sequencing using a multiple instance learning framework. Nat. Methods, 19, 1590–1598.

16. Begik, O., Lucas, M.C., Pryszcz, L.P., Ramirez, J.M., Medina, R., Milenkovic, I., Cruciani, S., Liu, H., Vieira, H.G.S., Sas-Chen, A., et al. (2021) Quantitative profiling of pseudouridylation dynamics in native RNAs with nanopore sequencing. Nat. Biotechnol., 10.1038/s41587-021-00915-6.

17. Kovaka, S., Hook, P.W., Jenike, K.M., Shivakumar, V., Morina, L.B., Razaghi, R., Timp, W. and Schatz, M.C. (2025) Uncalled4 improves nanopore DNA and RNA modification detection via fast and accurate signal alignment. Nat. Methods, 22, 681–691.

18. Maier, K.C., Gressel, S., Cramer, P. and Schwalb, B. (2020) Native molecule sequencing by nano-ID reveals synthesis and stability of RNA isoforms. Genome Res., 30, 1332–1344.

19. Vujaklija, I., Biđin, S., Volarić, M., Bakić, S., Li, Z., Foo, R., Liu, J. and Šikić, M. (2025) Detecting a wide range of epitranscriptomic modifications using a nanopore-sequencing-based computational approach with 1D score-clustering. Nucleic Acids Res., 53, gkae1168.

20. Huang, S., Zhang, W., Katanski, C.D., Dersh, D., Dai, Q., Lolans, K., Yewdell, J., Eren, A.M. and Pan, T. (2021) Interferon inducible pseudouridine modification in human mRNA by quantitative nanopore profiling. Genome Biol., 22, 330.

21. Cruciani, S., Delgado-Tejedor, A., Pryszcz, L.P., Medina, R., Llovera, L. and Novoa, E.M. (2025) De novo basecalling of RNA modifications at single molecule and nucleotide resolution. Genome Biol., 26, 38.

22. McCormick, C.A., Akeson, S., Tavakoli, S., Bloch, D., Klink, I.N., Jain, M. and Rouhanifard, S.H. (2023) Multicellular, IVT-derived, unmodified human transcriptome for nanopore-direct RNA analysis. bioRxiv, 10.1101/2023.04.06.535889.

23. Liu, H., Begik, O. and Novoa, E.M. (2021) EpiNano: Detection of m6A RNA Modifications Using Oxford Nanopore Direct RNA Sequencing. Methods Mol. Biol., 2298, 31–52.

24. Song, J., Lin, L.-A., Tang, C., Chen, C., Yang, Q., Zhang, D., Zhao, Y., Wei, H.-C., Linghu, K., Xu, Z., et al. (2025) DEMINERS enables clinical metagenomics and comparative transcriptomic analysis by increasing throughput and accuracy of nanopore direct RNA sequencing. Genome Biol., 26, 76.

25. Pryszcz, L.P., Diensthuber, G., Llovera, L., Medina, R., Delgado-Tejedor, A., Cozzuto, L., Ponomarenko, J. and Novoa, E.M. (2025) Rapid and accurate demultiplexing of direct RNA nanopore sequencing datasets with SeqTagger. Genome Res., 10.1101/gr.279290.124.

26. van der Toorn, W., Bohn, P., Liu-Wei, W., Olguin-Nava, M., Smyth, R.P. and von Kleist, M. (2024) Demultiplexing and barcode-specific adaptive sampling for nanopore direct RNA sequencing. bioRxiv, 10.1101/2024.07.22.604276.

27. Diensthuber, G., Pryszcz, L.P., Llovera, L., Lucas, M.C., Delgado-Tejedor, A., Cruciani, S., Roignant, J.-Y., Begik, O. and Novoa, E.M. (2024) Enhanced detection of RNA modifications and read mapping with high-accuracy nanopore RNA basecalling models. Genome Res., 10.1101/gr.278849.123.

28. Acera Mateos, P., J Sethi, A., Ravindran, A., Srivastava, A., Woodward, K., Mahmud, S., Kanchi, M., Guarnacci, M., Xu, J., W S Yuen, Z., et al. (2024) Prediction of m6A and m5C at single-molecule resolution reveals a transcriptome-wide co-occurrence of RNA modifications. Nat. Commun., 15, 3899.

29. Picardi, E., Fonzino, A., Fosso, B., Visci, G., Gissi, C. and Pesole, G. (2025) Ab initio detection of multiple epitranscriptomic modifications from ONT direct RNA sequencing data. Research Square, 10.21203/rs.3.rs-6343479/v1.

30. Cozzuto, L., Delgado-Tejedor, A., Hermoso Pulido, T., Novoa, E.M. and Ponomarenko, J. (2023) Nanopore Direct RNA Sequencing Data Processing and Analysis Using MasterOfPores. Methods Mol. Biol., 2624, 185–205.

31. Wurm, J.P., Griese, M., Bahr, U., Held, M., Heckel, A., Karas, M., Soppa, J. and Wöhnert, J. (2012) Identification of the enzyme responsible for N1-methylation of pseudouridine 54 in archaeal tRNAs. RNA (New York, N.Y.), 18.

32. Nance, K.D. and Meier, J.L. (2021) Modifications in an emergency: The role of N1-methylpseudouridine in COVID-19 vaccines. ACS Cent. Sci., 7, 748–756.

33. Gao, Y. and Fang, J. (2021) RNA 5-methylcytosine modification and its emerging role as an epitranscriptomic mark. RNA Biology, 18, 117.

34. Fleming, A.M., Mathewson, N.J., Howpay Manage, S.A. and Burrows, C.J. (2021) Nanopore dwell time analysis permits sequencing and conformational assignment of pseudouridine in SARS-CoV-2. ACS Cent. Sci., 7, 1707–1717.

35. Popova, A.M. and Williamson, J.R. (2014) Quantitative analysis of rRNA modifications using stable isotope labeling and mass spectrometry. J. Am. Chem. Soc., 136, 2058–2069.

36. Yang, J., Sharma, S., Watzinger, P., Hartmann, J.D., Kötter, P. and Entian, K.-D. (2016) Mapping of complete set of ribose and base modifications of yeast rRNA by RP-HPLC and mung bean nuclease assay. PLoS One, 11, e0168873.

37. Taoka, M., Nobe, Y., Yamaki, Y., Yamauchi, Y., Ishikawa, H., Takahashi, N., Nakayama, H. and Isobe, T. (2016) The complete chemical structure of Saccharomyces cerevisiae rRNA: partial pseudouridylation of U2345 in 25S rRNA by snoRNA snR9. Nucleic Acids Res., 44, 8951–8961.

38. Thüring, K., Schmid, K., Keller, P. and Helm, M. (2016) Analysis of RNA modifications by liquid chromatography-tandem mass spectrometry. Methods, 107, 48–56.

39. Taoka, M., Nobe, Y., Yamaki, Y., Sato, K., Ishikawa, H., Izumikawa, K., Yamauchi, Y., Hirota, K., Nakayama, H., Takahashi, N., et al. (2018) Landscape of the complete RNA chemical modifications in the human 80S ribosome. Nucleic Acids Res., 46, 9289–9298.

40. Zhang, Z., Chen, T., Chen, H.-X., Xie, Y.-Y., Chen, L.-Q., Zhao, Y.-L., Liu, B.-D., Jin, L., Zhang, W., Liu, C., et al. (2021) Systematic calibration of epitranscriptomic maps using a synthetic modification-free RNA library. Nat. Methods, 18, 1213–1222.

41. Makhamreh, A., Tavakoli, S., Fallahi, A., Kang, X., Gamper, H., Nabizadehmashhadtoroghi, M., Jain, M., Hou, Y.-M., Rouhanifard, S.H. and Wanunu, M. (2024) Nanopore signal deviations from pseudouridine modifications in RNA are sequence-specific: quantification requires dedicated synthetic controls. Sci. Rep., 14, 22457.

42. Baquero-Pérez, B., Yonchev, I.D., Delgado-Tejedor, A., Medina, R., Puig-Torrents, M., Sudbery, I., Begik, O., Wilson, S.A., Novoa, E.M. and Díez, J. (2024) N6-methyladenosine modification is not a general trait of viral RNA genomes. Nat. Commun., 15, 1964.

43. Cappannini, A., Ray, A., Purta, E., Mukherjee, S., Boccaletto, P., Moafinejad, S.N., Lechner, A., Barchet, C., Klaholz, B.P., Stefaniak, F., et al. (2024) MODOMICS: a database of RNA modifications and related information. 2023 update. Nucleic Acids Res., 52, D239–D244.

44. Delgado-Tejedor, A., Medina, R., Begik, O., Cozzuto, L., López, J., Blanco, S., Ponomarenko, J. and Novoa, E.M. (2024) Native RNA nanopore sequencing reveals antibiotic-induced loss of rRNA modifications in the A- and P-sites. Nat. Commun., 15, 10054.

45. Pryszcz, L.P. and Novoa, E.M. (2021) ModPhred: an integrative toolkit for the analysis and storage of nanopore sequencing DNA and RNA modification data. Bioinformatics, 38, 257–260.

46. Esfahani, N.G., Stein, A.J., Akeson, S., Tzadikario, T. and Jain, M. (2025) Evaluation of Nanopore direct RNA sequencing updates for modification detection. Genomics.

47. Hewel, C., Hofmann, F., Dietrich, V., Wierczeiko, A., Friedrich, J., Jenson, K., Mündnich, S., Diederich, S., Sys, S., Schartel, L., et al. (2024) Direct RNA sequencing (RNA004) allows for improved transcriptome assessment and near real-time tracking of methylation for medical applications. bioRxiv, 10.1101/2024.07.25.605188.

48. Guo, Z., Shao, Y., Tan, L., Lu, B., Deng, X., Chen, S. and Li, R. (2025) Enhanced detection of RNA modifications in Escherichia coli utilizing nanopore RNA004 technology. bioRxiv, 10.1101/2025.01.16.633309.

49. Rübsam, F.N.M., Liu-Wei, W., Sun, Y., Patel, B.I., van der Toorn, W., Piechotta, M., Dieterich, C., von Kleist, M. and Ehrenhofer-Murray, A.E. (2025) MoDorado: Enhanced detection of tRNA modifications in nanopore sequencing by off-label use of modification callers. Molecular Biology.

50. Zou, Y., Ahsan, M.U., Chan, J., Meng, W., Gao, S.-J., Huang, Y. and Wang, K. (2025) A Comparative Evaluation of Computational Models for RNA modification detection using Nanopore sequencing with RNA004 Chemistry. bioRxivorg, 10.1101/2025.02.03.636352.

51. Georgeson, J. and Schwartz, S. (2024) No evidence for ac4C within human mRNA upon data reassessment. Mol. Cell, 84, 1601–1610.e2.

52. Beiki, H., Sturgill, D., Arango, D., Relier, S., Schiffers, S. and Oberdoerffer, S. (2024) Detection of ac4C in human mRNA is preserved upon data reassessment. Mol. Cell, 84, 1611–1625.e3.

53. McCormick, C.A., Meseonznik, M., Qiu, Y., Fanari, O., Thomas, M., Liu, Y., Bloch, D., Klink, I.N., Jain, M., Wanunu, M., et al. (2024) mRNA psi profiling using nanopore DRS reveals cell type-specific pseudouridylation. bioRxivorg, 10.1101/2024.05.08.593203.

54. Sklias, A., Cruciani, S., Marchand, V., Spagnuolo, M., Lavergne, G., Bourguignon, V., Brambilla, A., Dreos, R., Marygold, S.J., Novoa, E.M., et al. (2024) Comprehensive map of ribosomal 2’-O-methylation and C/D box snoRNAs in Drosophila melanogaster. Nucleic Acids Res., 52, 2848–2864.

55. Baba, T., Ara, T., Hasegawa, M., Takai, Y., Okumura, Y., Baba, M., Datsenko, K.A., Tomita, M., Wanner, B.L. and Mori, H. (2006) Construction of Escherichia coli K-12 in-frame, single-gene knockout mutants: the Keio collection. Mol. Syst. Biol., 2, 2006.0008.

56. Cozzuto, L., Liu, H., Pryszcz, L.P., Pulido, T.H., Delgado-Tejedor, A., Ponomarenko, J. and Novoa, E.M. (2020) MasterOfPores: A Workflow for the Analysis of Oxford Nanopore Direct RNA Sequencing Datasets. Front. Genet., 11, 507358.

57. Li, H. (2018) Minimap2: pairwise alignment for nucleotide sequences. Bioinformatics, 34, 3094–3100.

58. De Coster, W. and Rademakers, R. (2023) NanoPack2: population-scale evaluation of long-read sequencing data. Bioinformatics, 39, btad311.

59. Bailey, T.L. (2021) STREME: accurate and versatile sequence motif discovery. Bioinformatics, 37, 2834–2840.

