## Supplemental_Figures for "Systematic benchmarking of basecalling models for RNA modification detection with highly-multiplexed nanopore sequencing"

### SUPPLEMENTARY FIGURES

**Figure S1. Benchmarking of a 96-barcoding model for the RNA004 chemistry. (A)** Confusion matrix of the 96 barcode mRNA model (b96_RNA004) generated on the validation data. Recorded precision is indicated on top. Zoomed panels of the 96x96 confusion matrix for eight barcodes are shown on the right. **(B)** Barplots comparing the computation time of b04_RNA004 and b96_RNA004, on a benchmarking dataset of 100.000 reads each. Bars represent the mean value with error bars indicating +/- 1 standard deviation. Dots represent individual replicates. Statistical significance was determined using a two-sided *t*-test (ns: $p > 0.05$, *: $p <= 0.05$, **: $p <= 0.01$, ***: $p <= 0.001$)**. (C)** BaseQ plots of 100.000 subsampled reads from a dropout sequencing run in which 93/96 barcodes were sequenced in a single flowcell. SCBC-21, SCBC-59, and SCBC-88 were skipped to determine cross-contamination. N represents the number of reads passing the default baseQ cutoff of >50. The value in brackets corresponds to the relative contribution of each barcode to the overall library.

**Figure S1** *(legend on previous page)*

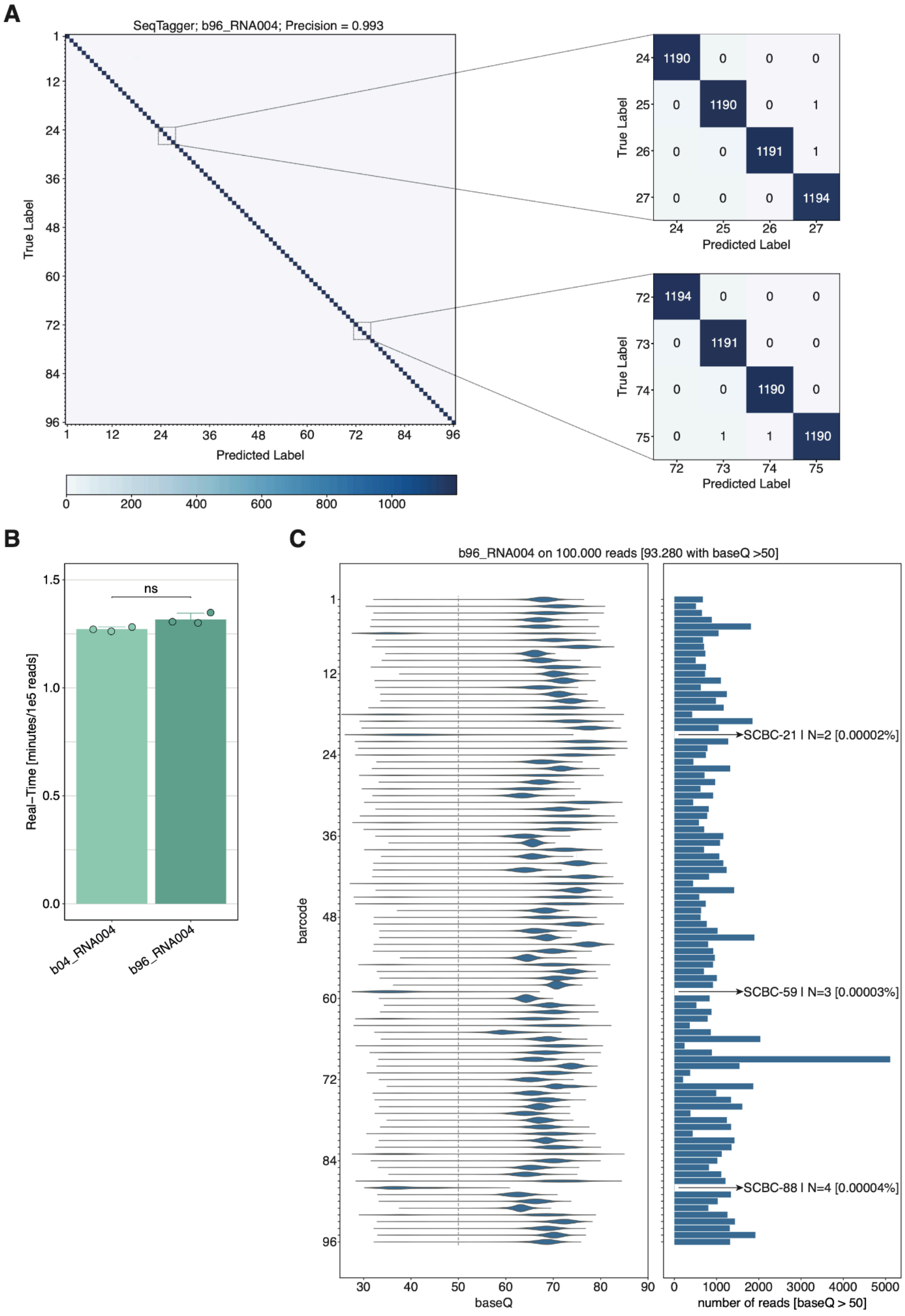

**Figure S2. Quality control of fully-modified and unmodified synthetic sequences from a highly multiplexed direct RNA sequencing run (A)** Barplots representing the overall uniquely aligned reads per replicate for each modification type (UNM, $m^6A$, Ψ, $m^1Ψ$, $m^5U$, $m^5C$, $hm^5C$, and $ac^4C$) across all four curlcake sequences (CC1-4) and basecalled with two accuracy modes (*hac* and *sup*). **(B)** 2D-density plots of the cumulative read quality and read length across the entire sequencing run, for both accuracy modes (*hac* and *sup*).

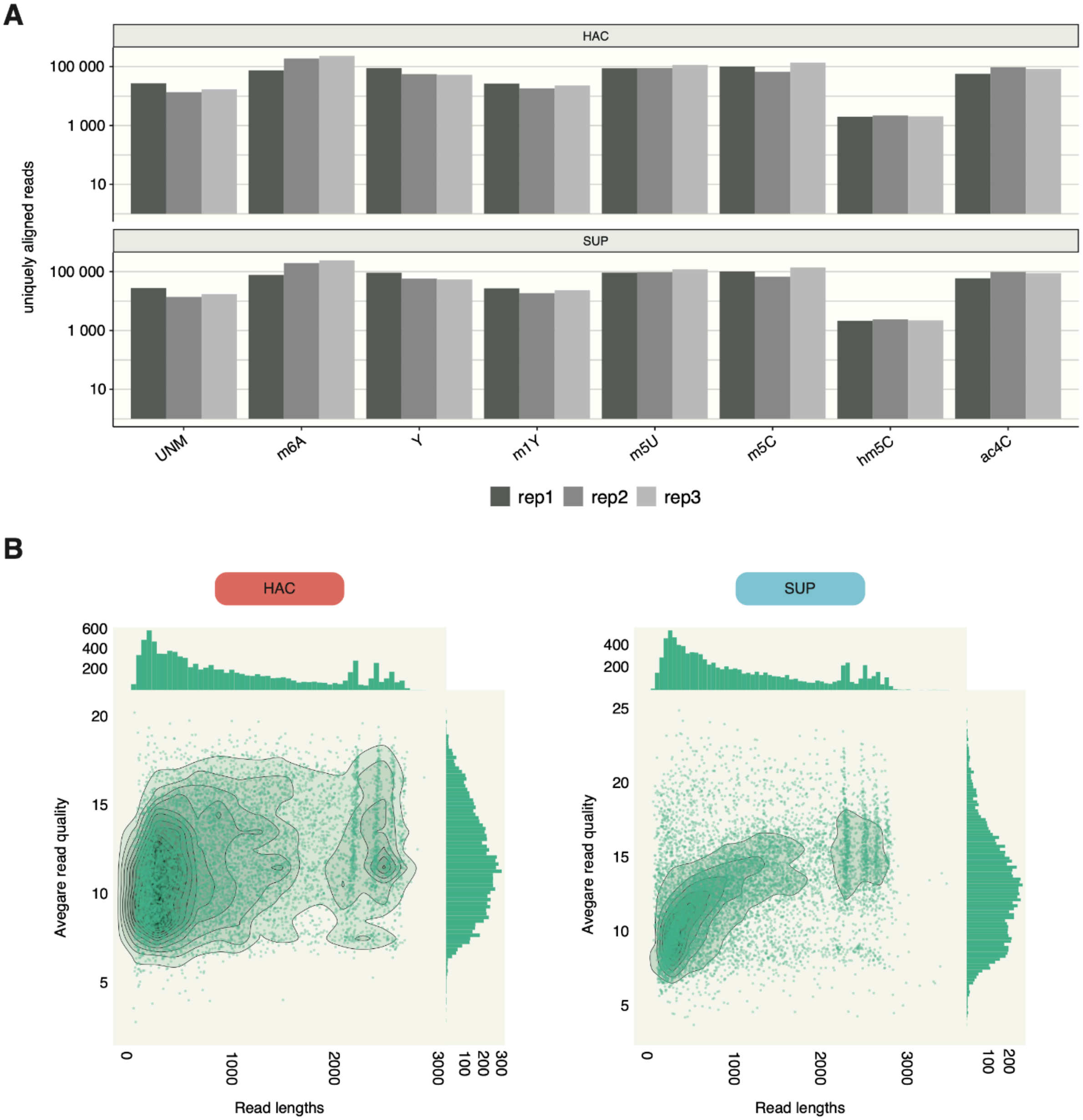

**Figure S3. Systematic assessment of the accuracy and cross-reactivity of pre-trained modification-aware basecalling models across a panel of synthetic sequences with diverse RNA modifications. (A)** Per-read modification probabilities reported for each model on fully modified and unmodified sequences. The abline represents the median modified frequency (expected = 100%). **(B)** ROC curves based on modification probabilities per read and site for each model. **(C)** Precision-Recall (PR) curve based on modification probabilities per read and site, for each basecalling model. The represented motifs cover the following sequences: KGACY (K = G/T,  and Y = C/T), BBABB (B = C/G/T), VVTVV (V = C/G/A), and DDCDD (D = A/G/T).

**Figure S3** (legend on previous page)

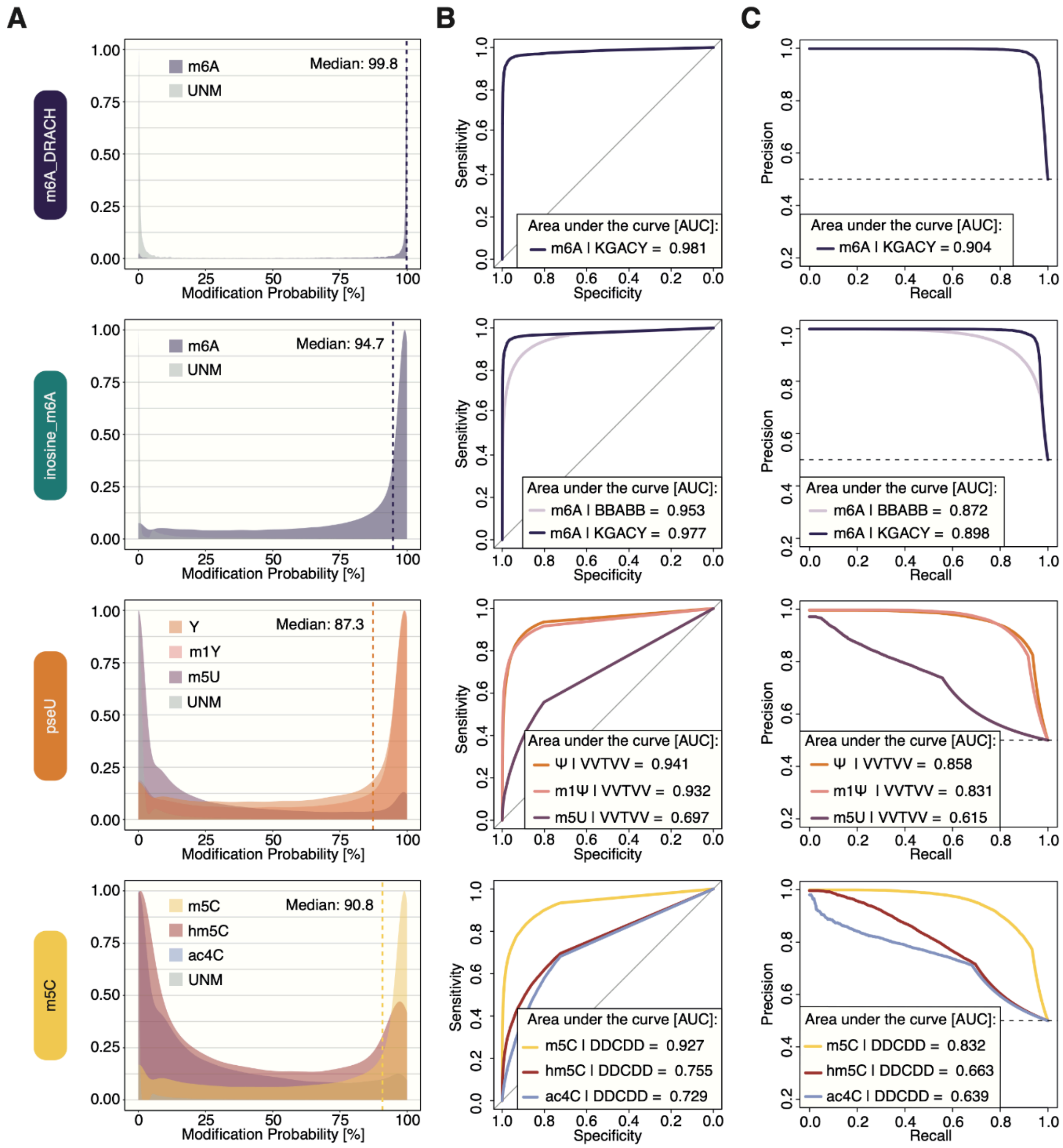

**Figure S4. Performance of pseU (A) and m5C (B) modification-aware *dorado* basecalling models in rRNA molecules**. In the upper panels, precision-recall (PR) curves are shown, both for *S. cerevisiae* and *E. coli* rRNAs using either of the two models. In the lower panels, Receiver Operating Characteristic (ROC) curves are shown for rRNA molecules from *S. cerevisiae* (left) and *E. coli* (right), using either of the two models. The Area Under the Curve (AUC) for PR and ROC curves is shown for each sample and basecalling model.

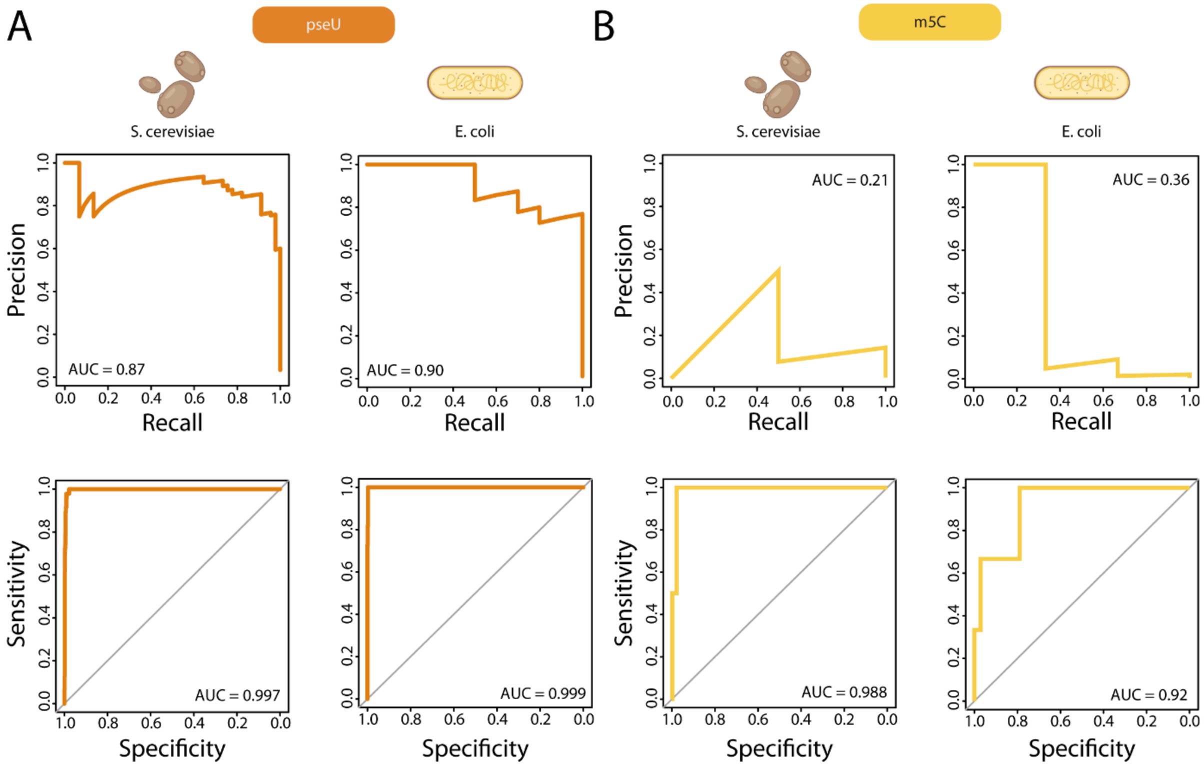

**Figure S5. Correlation of modified sites identified across different species for wildtype (WT) and *in vitro* transcribed (IVT) samples**. **(A)** Comparison of modified sites identified by WT (top) and IVT (bottom) using the pseU model. Pearson's correlation is indicated at the top left corner. **(B)** Comparison of modified sites identified by WT (top) and IVT (bottom) using the m5C model. Pearson's correlation is indicated at the top left corner.

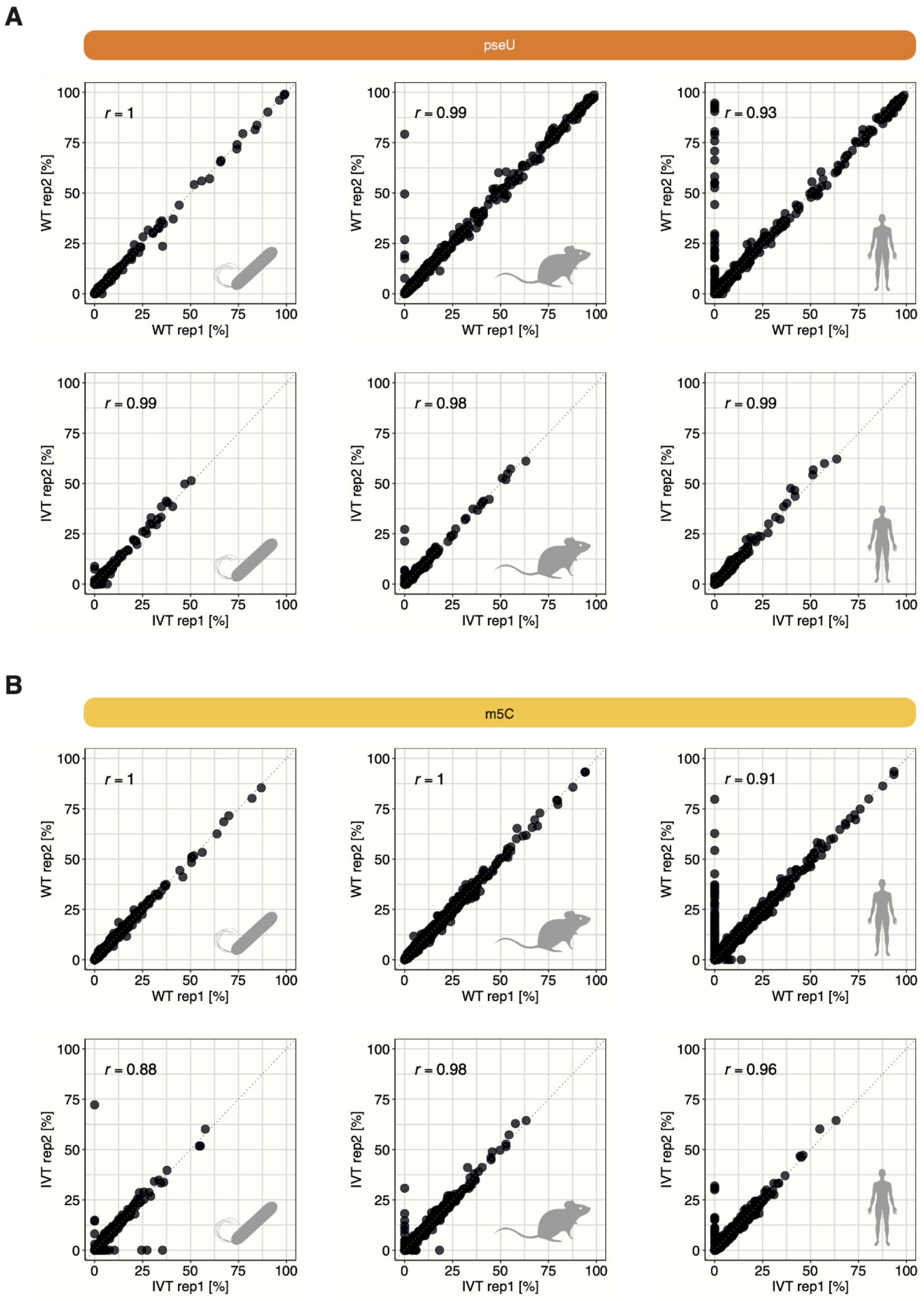

**Figure S6. Predicted modification stoichiometries for annotated modifications found on the two major rRNAs of *E. coli, M. musculus, and H. sapiens.* (A)** *Top:* Stoichiometry predictions of the pseU model for individual modifications found on the two major rRNAs of *E. coli, M. musculus, and H. sapiens*. *Bottom*: Stoichiometry predictions of the pseU model split into different categories, including non-targeted modifications (U_mods_non_Y) and predicted sites containing modified positions at the +/- 1nt position. **(B)** *Top:* Stoichiometry predictions of the m5C model for individual modifications found on the two major rRNAs of *E. coli, M. musculus, and H. sapiens*. *Bottom*: Stoichiometry predictions of the pseU model split into different categories, including non-targeted modifications (C_mods_non_Y) and predicted sites containing modified positions at the +/- 1nt position.

**Figure S6** (legend on previous page)

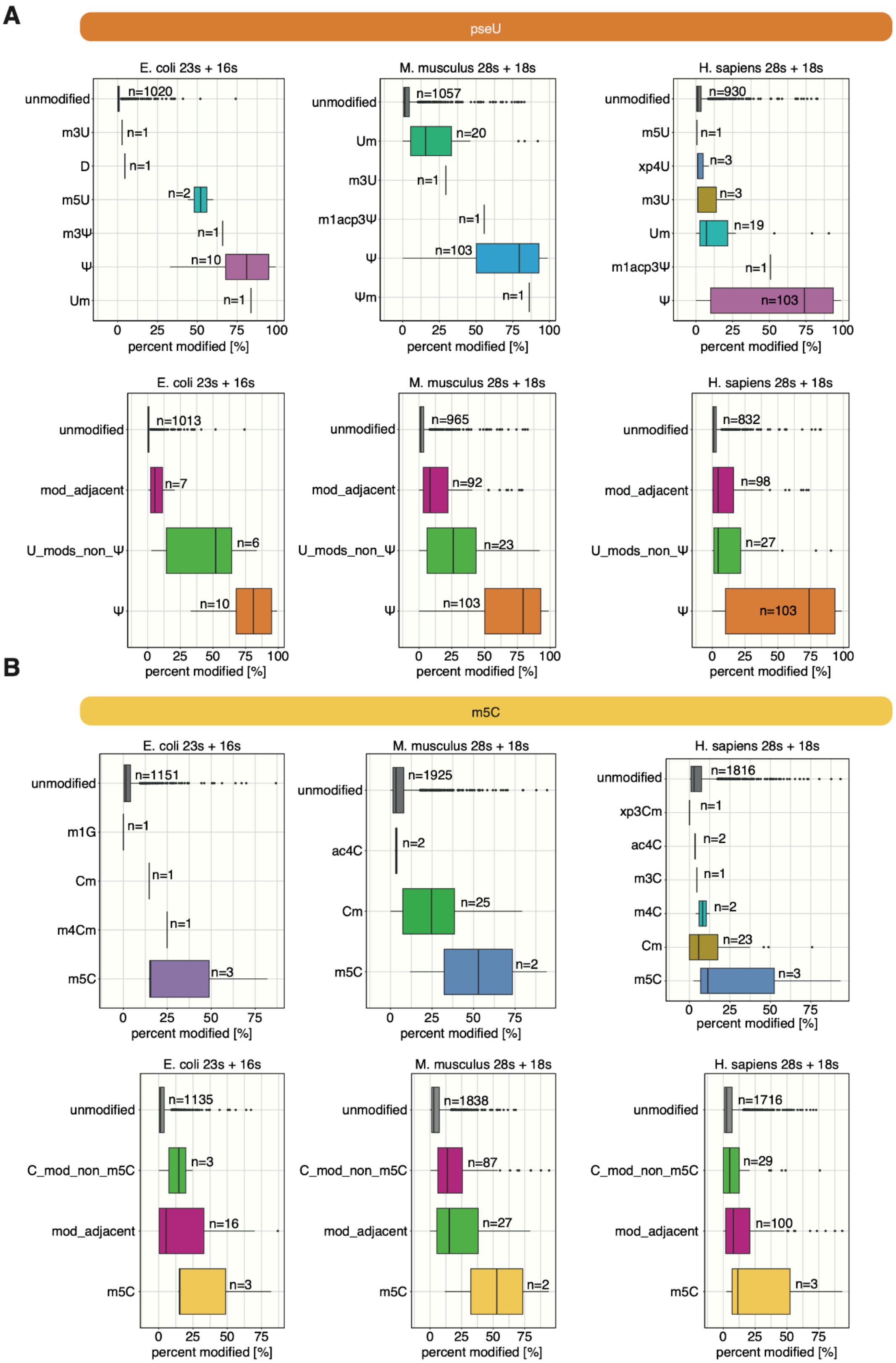
